# ProGuide: a flexible framework for modeling global conformational rearrangements in proteins using DEER-derived distance restraints

**DOI:** 10.1101/2025.06.05.658127

**Authors:** Julian D. Grosskopf, Peter Kasson, Michael T. Lerch

**Affiliations:** Department of Biophysics, Medical College of Wisconsin, Milwaukee, WI 53226, USA; Departments of Chemistry and Biochemistry and of Biomedical Engineering, Georgia Institute of Technology, Atlanta, GA 30332 USA

## Abstract

Conformational heterogeneity is integral to protein function – ranging from enzyme catalysis to signal transduction – and visualizing distinct conformational states requires experimental techniques capable of providing such structural information. One particularly powerful method, double electron-electron resonance (DEER) spectroscopy, can provide a high-resolution, long-range (∼15-80 Å) probability distributions of distances between site-selected pairs of spin labels to resolve intra-protein distance parameters of unique protein conformations, as well as their respective likelihoods within a conformational ensemble. A current frontier in the field of DEER spectroscopy is utilizing this distance information in computational modeling to generate complete structural models of these multiple conformations. Although several methods have been developed for this purpose, modeling protein backbone structural rearrangements using multiple distance restraints remains challenging, due in part to the complexity provided by rotameric flexibility of the spin label side chain. Here, we overcome these challenges with ProGuide, a new framework for generating accurate structural models guided by DEER distance distribution information. Large conformational rearrangements are captured by performing iterative experimentally biased molecular dynamics simulations. In each iteration, spin-label rotameric heterogeneity is modeled using chiLife, and then Cα changes are calculated to capture distance-probability density present in the experimental DEER distributions and lacking from the modeled one. The resulting models of this process then go through a selection to generate the ensemble that best recapitulates the DEER data. We illustrate the power of this method using published DEER data from a study of biased agonism in the angiotensin II type 1 receptor (AT1R), a prototypical G protein coupled receptor (GPCR). The resulting AT1R models consist of both Gq- and β-arrestin-biased conformations, including a completely novel β-arrestin-biased conformation. These models reveal structural insights involving tertiary structural rearrangements as well as residue-level changes in crucial microswitch motifs. Taken together, the results demonstrate the power and flexibility of ProGuide to investigate conformational rearrangements of large, complex proteins using DEER-derived distance restraints.

## Introduction

Proteins are dynamic molecules that exhibit fluctuations in structure across a wide range of time and length scales. A hierarchical view of protein dynamics is encoded by an energy landscape perspective of protein conformational heterogeneity^1–4^, in which the relative populations and lifetimes of distinct states are dictated by their equilibrium free energies and the energy barriers between states, respectively ^5–7^. Unique conformational states are often distinguished by collective protein backbone motions such as changes in secondary or tertiary structure on the microsecond-to-millisecond timescale. Each conformational state is comprised of multiple statistical sub-states with picosecond-to nanosecond lifetimes that exhibit low-amplitude (free energy barrier < ∼10 kT) deviations in backbone and sidechain position from the median structure. Modulation of the energy landscape (i.e., changes in conformational equilibria or lifetimes) may be driven by a variety of factors including changes to the cellular environment, post-translational modifications, or interaction with binding partners such as small molecule ligands or other proteins. This modulation, in turn, governs cellular processes such as enzyme catalysis, import/export processes, cellular signaling pathways, and many others.^8^ Defining the ensemble of conformations adopted by a protein in solution is crucial to uncovering molecular mechanisms that underly normal cellular processes, disease-associated dysregulation, and understanding/discovering therapeutic interventions of these mechanisms.

Mapping a protein’s conformational energy landscape requires an integrated approach combining structural, biophysical, and computational tools. High-resolution atomic structures of individual conformations are accessible through structural biology methods such as X-ray crystallography and cryo-EM. These techniques typically capture the most stable (i.e., lowest free energy) conformational states of a protein, and significant effort is often required to stabilize and resolve alternative conformations. Despite recent breakthroughs in image processing^9–11^ and multi-temperature^12–14^ and time-resolved measurements^15,16^, it remains challenging to capture the full complexity of the conformational ensemble with cryo-EM and X-ray crystallography. To this end, complementary spectroscopic techniques such as NMR, FRET, and EPR have proven extremely valuable in revealing more sparsely populated or transient conformations and monitoring more flexible protein regions^17–21^.

Over the last two decades, the pulsed EPR experiment DEER (double electron-electron resonance)^22^ paired with site-directed spin labeling (SDSL) has emerged as a crucial spectroscopic technique for monitoring protein conformational heterogeneity. In SDSL, site-specific spin labels are attached to two residues on the protein by means of directed cysteine mutagenesis and addition of a thiol-reactive spin label^23^. Fitting the DEER spin-echo decay then reports distances between the spin labels in the 15-80 Å range with sub-angstrom precision. Importantly, DEER yields a probability distribution of distances between spin labels, rather than an average distance, enabling the detection of each conformation present in the ensemble as well as providing their relative populations. An individual DEER experiment provides sparse information about the global conformation of a protein, but technological advances have made it possible to collect increasingly larger DEER datasets consisting of multiple spin pairs in various conditions (e.g., different ligands), from which correlated conformational changes may be identified. The strength and utility of DEER to probe conformational rearrangements of globular and membrane proteins has been detailed in numerous reviews^22,24,25^ and studies revealing mechanistic insight into signal transduction^17,18,26–29^, ion transport^30^, and antibiotic resistance^31,32^, among others. Importantly, these studies establish the role DEER plays in discovering structure-based functional mechanisms as they relate to other functional experiments. One of the limitations of DEER is that distance distributions provide information that can be used to *infer* conformational rearrangements with respect to a reference structural model, but an atomistic model cannot be derived from DEER data alone. The potential for combining DEER and computational modeling to overcome this limitation is well-recognized^33,34^, and using DEER results as restraints in computational modeling is a current frontier in the field^35–40^. The current state of the field, including limitations of existing approaches, are discussed in detail in a pair of excellent recent reviews^33,34^.

The different methods available for integrating DEER data into computational modeling can be broadly categorized as physics-based (e.g., molecular dynamics (MD) simulations, Rosetta) or statistics-based (e.g., Alphafold3, Alphafold2, RoseTTAFOLD2). Deep learning approaches such as Alphafold use statistical-based methods to rapidly generate large sets of conformations, which can then be reweighted based on their consistency with experimental DEER data.^28,41^ Physics-based methods such as MD simulation can be used to refine deep-learning models and include chemical information such as parameters for bond length, bond geometries, atomic interaction potentials, and electrostatics^42^.

The spin-label side chain poses a major challenge to integrating DEER data in MD simulations. Biased MD simulation methods using explicit spin labels can significantly alter the conformation of the spin-label nitroxide group, producing inaccurate spin label rotameric ensembles. The resulting protein conformations thus may be unphysical.^34^ Explicit spin labels also require parameterization of the spin label side chain, which is not trivial and poses a barrier for refinement of data from promising new labels ^43–45^ as well as routinely used spin labels (e.g., IAP^18,46,47^ and V1^26,48,49^) that have not yet been parameterized. Alternatively, some methods omit the spin label altogether, using experimental distance distributions between nitroxides as distance restraints between C_α_ residues ^40^, which can lead to nanometer-scale uncertainties^37^.

In the present work, we introduce ProGuide, a framework to overcomes these limitations, enabling the modeling of both backbone and spin-label conformational rearrangements using multiple distance restraints. ProGuide sidesteps the explicit spin label challenge by separating calculation of spin label rotamers from calculation of protein backbone changes. To achieve the first, it leverages the flexible and powerful spin label rotamer library chiLife^50^. To achieve the second, it utilizes iterative biased molecular dynamics (MD) simulations built upon the bias-resampling ensemble refinement (BRER)^40^ engine, where a corrective bias potential is applied based on the difference between the simulated rotamer distribution and an experimental target. ProGuide facilitates the use of experimental distance restraints generated with any spin label to drive backbone rearrangements in the modeled protein, achieving excellent overlap between modeled and experimental distance distributions.

We demonstrate the utility of ProGuide in applications to a pair of well-studied G protein coupled receptors (GPCRs): the β_2_ adrenergic receptor (β2AR) and the angiotensin II type I receptor (AT1R). GPCRs are the largest family of membrane proteins in humans and serve as the primary conduit for transmitting extracellular signals into the cell. This typically occurs through binding of a diffusible ligand at the extracellular surface of the receptor, regulating receptor activation of various intracellular proteins including heterotrimeric G proteins and β-arrestins^51–55^. Ligands may induce preferential engagement of specific G proteins or β arrestins, a phenomenon known as biased signaling. The prevailing model indicates that ligand bias is achieved through stabilization of distinct conformations or conformational equilibria of the receptor^26,49,56^, with some of the strongest evidence for this mechanism coming from DEER studies of the AT1R^26^.

DEER has played a pivotal role in uncovering molecular mechanisms of function in several GPCRs, including the AT1R and β2AR^17,18,26,27,29,57^. As an initial demonstration of ProGuide, we model the transition of transmembrane helix 6 (TM6) of the β2AR from an inactive inward position to an active outward position (∼10 Å helical rigid-body displacement) using DEER data collected on a single spin label pair of the β2AR (Figure 1A). Then, to demonstrate the capability of ProGuide to model coordinated and complex rearrangements, we incorporated multiple experimental datasets simultaneously and generated models of global conformational rearrangements of the AT1R. Specifically, we used DEER data previously collected on 10 different spin label pairs of the AT1R (Figure 1B) to model unique conformations linked to signaling bias^26^. The AT1R models generated here reveal distinct tertiary structural rearrangements and statistically significant changes in conserved receptor motifs, including identification of a previously unreported conformation implicated in β arrestin biased signaling. Identification of these conformational details highlights the advantage of obtaining complete atomistic models from experimental DEER data.

**Figure 1:**
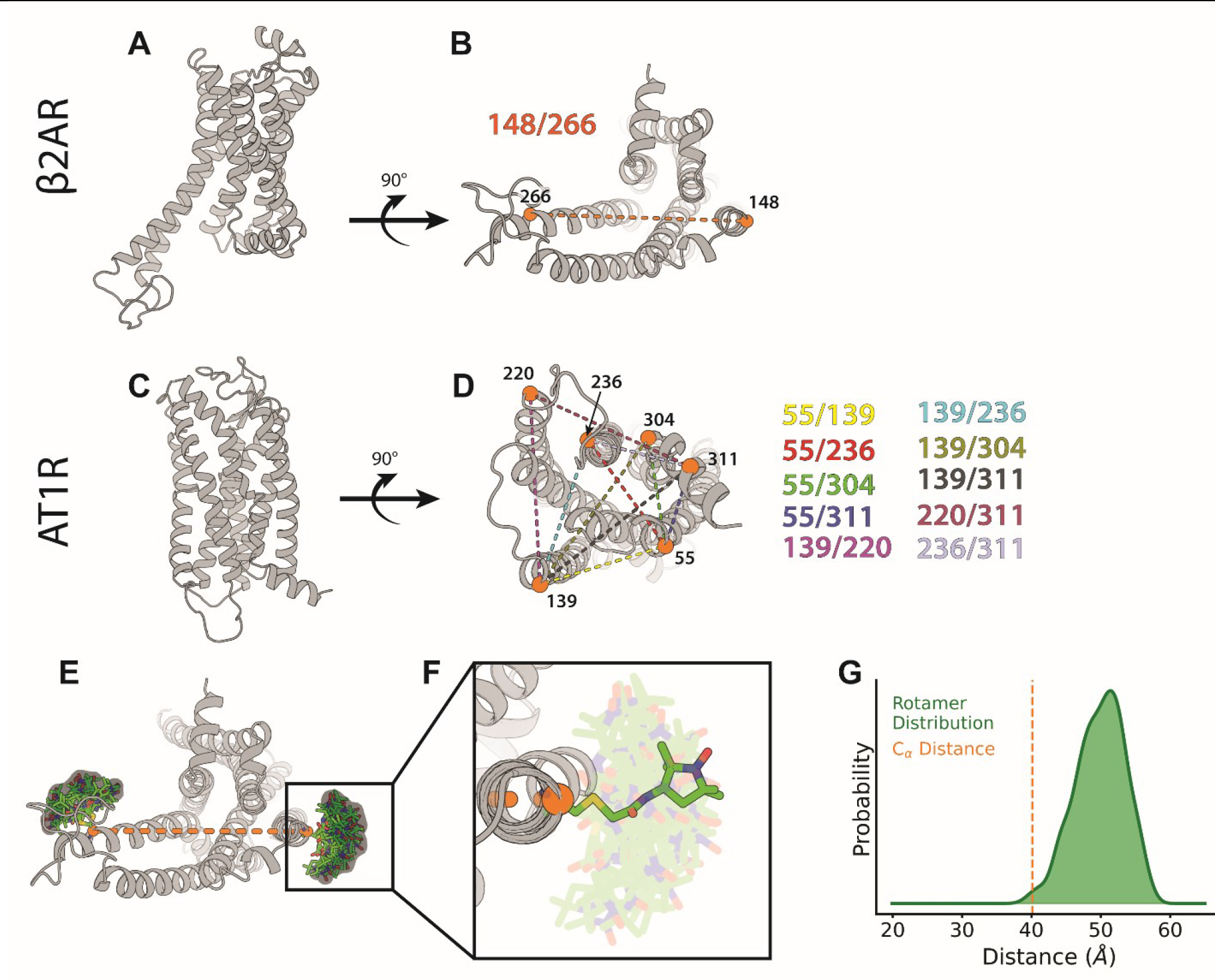
Label sites and spin-label rotameric heterogeneity. A) and B) show the β2AR viewed from the membrane plane and the cytoplasmic surface, respectively, with the spin label site Cα atoms shown as orange spheres. C) and D) show the AT1R viewed from the membrane plane and the cytoplasmic surface, respectively, with the spin label site Cα atoms shown as orange spheres. E) demonstrates a superimposed rendering of IAP rotamers (green) modeled on to the β2AR (grey cartoon, PDB ID: 3SN6, loops modelled in), shown from the cytoplasmic face. Transparent clouds represent the nitroxide N-O atoms where the free radical is trapped, which ultimately serves as the point of measurement in DEER spectroscopy. Orange spheres represent the C_α_ atoms that harbor the spin label, with an orange dashed line signifying the distance between C_α_ atoms. F) shows a magnified view of the IAP spin label, with a single rotamer highlighted. G) illustrates a graph of the IAP nitroxide distribution (green) and distance between C_α_ atoms that harbor the IAP spin label (orange dotted line).

## Methods

ProGuide incorporates the BRER simulation engine^40^ and chiLife^50^ with the iterative modeling by residual space exploration (IMRSE) algorithm, allowing for the use of multiple experimental distance distributions to drive large and complex global conformational rearrangements. To begin this process, an input structural model is selected or created (experimental structure from crystallography/cryo-EM/NMR or computational from AlphaFold or similar). ChiLife is used to model spin label side-chain rotamers on the protein and create a simulated distance distribution for each spin-pair used in DEER experiments (**Figure 1, Figure 2A**). Probability-distance distributions are specified by the following:

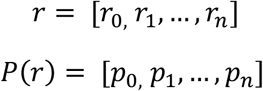

**Figure 2:**
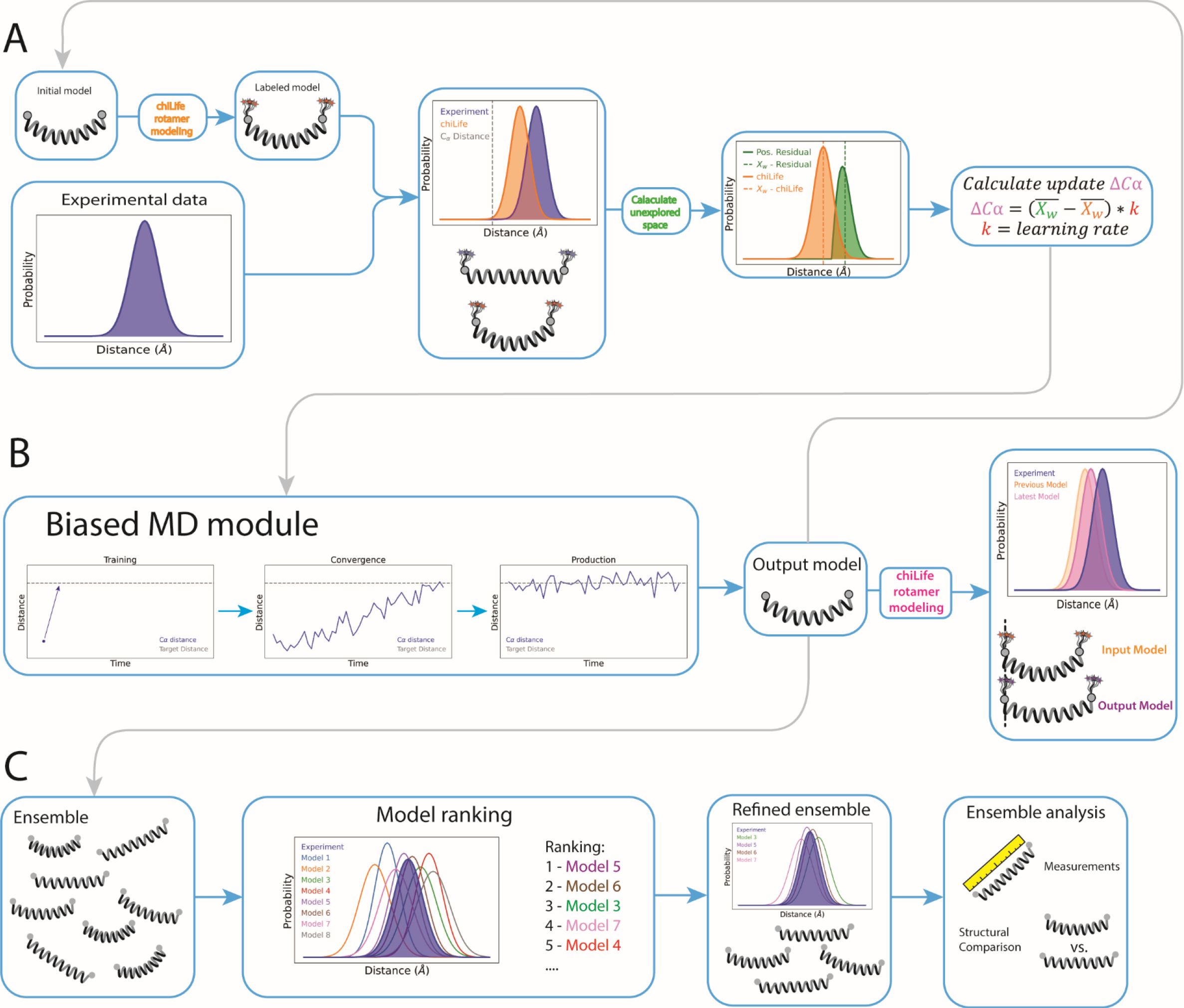
Schematic of ProGuide. A) Update block. This block calculates an incremental update to the Cα distance between the two spin-labeled residues. First, an input model and experimental data are fed into ProGuide. Using chiLife, the simulated distribution between the two spin-labeled residues is calculated from the coordinates of the input model (orange distribution). The simulated distribution is compared with the experimental data and the unexplored space of the experimental distribution is calculated by subtracting the simulated distribution from the experimental data and keeping only the positive residuals (green). An update to the Cα distance is then calculated (see methods) and applied to the next block in the framework. B) biased MD block. This block receives the updated Cα distance and performs biased MD in 3 phases: training phase calculates a linear bias potential to drive the necessary change; convergence phase applies the bias potential to the system and ends once the distance between Cα atoms in the simulation is within a certain cut off (typically 1-2 Å); production phase allows the system to relax around this bias potential for some amount of time. Using chiLife, the last frame of the production phase is then used to simulate the spin-label distribution that this model provides. An example of the simulated distribution from the output model compared to the input model and experimental data is shown. The output model is then fed back into the update block to calculate a new update to the Cα distance, as well as fed into the final block. C) Model ranking and analysis block. The ensemble of output models is reweighted (the component models can also be ranked) to generate spin label rotamer set/distribution that best overlays with the experimental data. The top *n* model coordinates can be output for further structural and statistical analysis.

Where *r* is the distance vector and *P*(*r*) is the probability vector as a function of distance. The simulated distribution is constructed such that the distance vector (*r*) is identical to the experimental distance vector to allow for point-wise calculations between the distributions. The simulated distribution is subtracted from the experimental distribution, and a positive residual function (PRes) is applied to the resulting residual distribution vector:

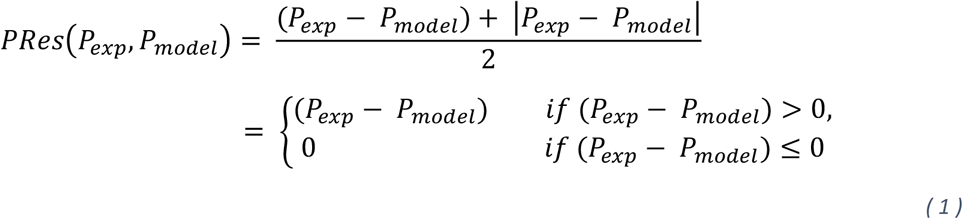

where *P*_*exp*_ is the probability vector of the experimental probability-distance distribution and *P*_*model*_ is the probability vector of the simulated probability-distance distribution from the input model protein. The PRes function returns all positive values and replaces negative values with zero, thus identifying all areas of the experimental distribution that are not sampled by the input model distribution. This is referred to as the positive residual distribution. A momentum parameter is also specified to control how many previous models to incorporate into the positive residual distribution calculation as follows:

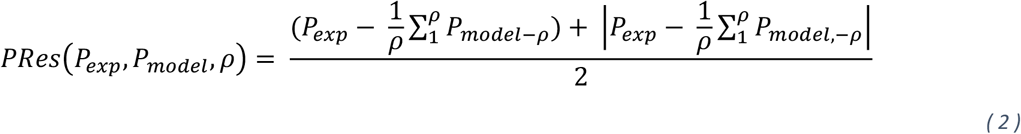

where ρ indicates the momentum parameter and *P*_*model*−1_ is the probability vector from the simulated probability-distance distribution of the latest generated model, *P*_*model*−2_, is the model generated prior to *P*_*model*−1_, and so on. The 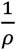 term is a normalization term, assuming *P*_*exp*_ and *P*_*model*_ are already area normalized to one another. A value greater than 1 specifies multiple previous models and is useful in cases wherein the width of the input model distribution is narrower than the experimental distribution so that the entirety of the experimental distribution is explored. To visualize the impact of ρ values on distribution convergence, see supplemental information and Figure S9.

The probability weighted average distance of a probability distance distribution can be calculated as follows:

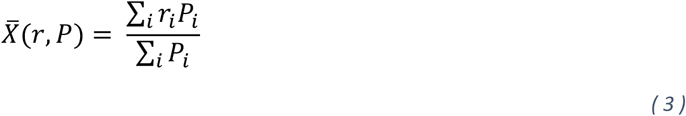

To determine the target distance used in MD simulations, an update to the Cα distance (Δ*Cα*) is calculated according to equation 1,

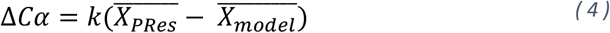

Where 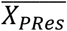 and 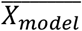 is the probability weighted average distance of the PRes and model distributions, respectively, and the update rate *k* is a value between 0 and 1 used to control the size of the update applied. In this application, a *k* value lower than 1 reduces the magnitude of the change in distance between Cα atoms imposed in biasing simulations, providing for relatively small perturbations to the model. The effects of varying *k* values on the model system is discussed in the supplemental information and visualized in Figure S9. Δ*Cα* is then added to the distance between Cα atoms of spin-labeled sites in the input model to produce the target distance, specified as:

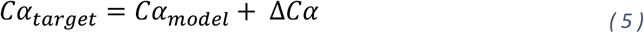

The target distance *Cα*_*target*_ and input model are fed into the BRER biased MD engine. The BRER engine follows three phases to apply biasing potentials to match experimentally derived distance values: training, convergence, and production phases. The training phase determines a linear bias potential needed to drive the Cα-Cα distances to the targets determined in the Cα update step. Importantly, in the case of multiple experimental restraints, linear bias potentials for all restraints are optimized simultaneously during the training phase. Once training is completed for a particular restraint, the bias potential is extracted, and training continues for the remaining restraints. The convergence phase applies the learned bias potentials to the system until Cα distances converge to the target distances, at which point the system is taken to the production phase. In the production phase, the simulation continues for a user-specified amount of time with bias potentials intact, which allows the system to relax from the abrupt structural transition induced during convergence. The last frame in the production phase is used as an “output model” (**Figure 2B**), which is then used as the input model for the next iteration of model generation.

To summarize the model generation process of ProGuide, the output model of the latest iteration is used as the input model to generate new distance targets for generation of the next model. Then, the new input model and new distance targets are fed into the BRER engine and bias potentials are recalculated according to the new targets. Upon completion of the biased simulations, the output model is used as the new input and the process begins again. This is repeated for a user-specified *n* iterations, successively using the output model of the latest round of model generation as input for the next round (**Figures 2A and B**). Ten frames from the last ns of the production phase (one every 0.1 ns) from each iteration are collected to increase the sample size used in the analysis block (**Figure 2C**).

To select representative models that agree with the experimental data, each model distribution is ranked against the experimental data by means of the Jensen-Shannon (JS) divergence or inverse dot product. The JS divergence is calculated using equation 6,

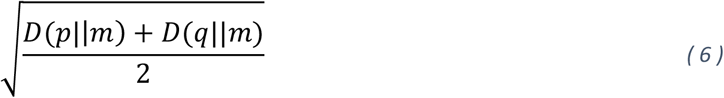

where *p* and *q* are the modeled and experimental distributions, *D* is the Kullback-Leibler divergence, and *m* is the point-wise mean of distributions *p* and *q*. Alternatively, similarity between modeled and experimental distributions can be calculated via the inverse dot product,

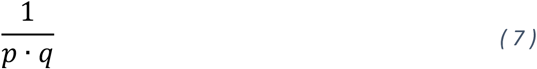

where the magnitude of the dot product is proportional to the similarity of the two vectors. A smaller value for JS divergence or inverse dot product indicates greater similarity between model and experimental distributions.

The linear combination of the top *n* ranked models represents a final ensemble that is compared to the experimental distributions using JS divergence. In the case of the AT1R, models were ranked by the inverse dot product (equation 7) where a lower value indicates better agreement with the experimental data. This was found to produce a smaller JS divergence in the top 50 summed models compared to the experimental data. For the β2AR, ranking models using the JS divergence resulted in a final ensemble that best agreed with the experimental data.

### BRER

BRER^40^ is a parallel, iterative ensemble refinement method that utilizes biased MD simulations implemented in GROMACS^58^ 2019.6 using gmxapi^59^. Briefly, BRER creates new models by means of a three-phase process: training, convergence, and production. A learned maximum-entropy linear bias potential ^60,61^ (training phase) is applied to the whole system and drives two atoms to a target distance (convergence phase), after which the applied linear bias is maintained for a user-specified period of time to allow the rest of the system to relax (production phase).

Here, instead of using the bias-resampling ensemble refinement that forms the BRER algorithm, we eliminate the probability weighted draw from the input distribution and instead utilize a single target distance between each atom pair. This utilizes the linear potential framework implemented in BRER but not its high-level algorithm for refinement. Identification of target distances is described in *Method Overview* and depicted in **Figure 2A**. The system inputs required here are a starting MD system and target distances.

Primary adjustable parameters include *A, tolerance*, and τ (tau). *A* is the learning rate for the coupling constant α (i.e., bias potential). The *A* value is a critical parameter for determining convergence and behavior of the system. *A* values set too small will not enable proper convergence on the target distance, and values set too large may produce erroneous models due to a highly dissipative convergence process. The ideal value of *A* is the maximum value of α expected for the given transformation. If a larger coupling constant is needed, the program will search above the set *A* value in the training phase until a suitable α is found. For more information on the A parameter, we refer the interested reader to the following articles^40,60^. *Tolerance* specifies how close the distance between atoms has to be to the specified target distance, in nm, before the simulation moves from the convergence to the production phase. The τ value describes the length of time in ps the system is allowed to adapt to the new α in the training phase before either re-adjusting the α value or accepting the current value and moving to the convergence phase. The τ value can have an impact on the learned α, where higher values of τ introduce lower coupling constant values. The parameters of *A, tolerance*, and τ used in this work are provided in **Table S1**.

BRER is maintained by the Kasson Lab and is available on GitHub (https://github.com/kassonlab/run_brer).

### ChiLife

ChiLife^50^ is an open-source Python package used to simulate distance distributions of EPR spin labels attached to protein structures/models *in-silico*, and is highly amenable to unique spin labels, rotamer libraries, and rotamer sampling methods. For the IAP label, used herein on the β2AR, the rotamer library was expanded through a modified version of off-rotamer sampling in which spin label side chain dihedral angles were randomly altered using a standard deviation of 15° from the underlying rotamer library. For the V1 label used on the AT1R, the rotamer library was expanded in the same manner with dihedral angle standard deviation of 10°. The dihedral angle standard deviation values specified above were the minimal values that generated reproducible *in-silico* labeling and distance distributions. The rotamer sets created using this approach were used for their respective proteins through the entirety of model generation and analysis.

ChiLife is maintained by the Stoll Lab and is available on Github (https://github.com/StollLab/chiLife).

### MD system setup for modelling

In all cases, CHARMM-GUI^62^ was used to set up the initial system, with missing loop regions modelled in using Modeller^63^ in ChimeraX^64^. For the β2AR, the carazolol- and inactive-state stabilizing nanobody NB60-bound crystal structure was used as the starting model (PDB ID 5JQH)^65^. The N terminal T4 lysozyme addition, NB60 nanobody, and carazolol were removed. Mutations consistent with the β2AR construct used in the DEER experiments^18^ were introduced: C77V, C265A, C327S, C341L, C378A, C406A, M96T, and M98T. Titratable residue protonation assignments were determined at pH 7, with the exception of the following residues which were protonated consistent with previous studies^66^: E107, E122, D130, H172, E268. Disulfide bonds were specified between residues C184/C190 and C106/C191. Box dimensions were set to 90 Å x 90 Å with 22.5 Å above and below the bilayer composed of 201 POPC lipids and solvated with 20,358 water molecules. 0.15 M NaCl was added under neutralizing conditions for a total of 55 Na and 66 Cl ions.

For the AT1R, the 4YAY^67^ crystal structure was used as the starting model with the extracellular BRIL truncated. The conditions used to set up this system were consistent with those present in the Wingler et. al, 2019 MD simulations ^26^. Hydrogens were added to the structure and titratable residues were protonated in accordance with their dominant protonation state at pH 7. Residues D74 and D125 were deprotonated, consistent with observations in other GPCRs^68^. N and C termini capping was added with acetyl (ACE) and methylamide (NT3) groups, respectively. Disulfide bonds were specified between residues C101/C180 and C18/C274. An 80 × 80Å box was created composed of 153 POPC lipids and solvated with 85 water molecules per lipid for a total of 13,005 water molecules. 0.15 M NaCl was added under neutralizing conditions for a total of 41 Na and 55 Cl ions.

GROMACS^69^ (2019.6) was used for all MD simulations, and CHARMM36m^70^ was used as the forcefield in all simulations with a TIP3P water model. Relaxation using steepest descents was performed to a tolerance of 1.195 kcal/mol/Å^2^. Positional restraints were set at 10 kcal/mol/Å^2^ and an initial equilibration was performed for 20 ns using a 1 fs timestep. For the β2AR and AT1R, restraints were decreased to 5 kcal/mol/Å^2^ and tapered off across 60 ns and 80 ns, respectively, using NPT conditions with a velocity rescaling thermostat^71^ and a C-rescale barostat^72^ and a leapfrog integrator with a 2 fs timestep. Restraints were removed and each system was allowed to equilibrate for 50 ns in which the starting model for training and application of bias potentials was selected. All hydrogen atom bond lengths were constrained using the LINCS algorithm. All short-range electrostatics and Van der Waals interactions were calculated with a 12 Å cutoff and switching at 10 Å. All long-range electrostatic interactions were computed with the Particle-Mesh-Ewald method^73^.

### Model filtering, ranking, and selection

After model creation, spin label side chain rotamers were simulated five times for each spin pair, and the resulting modeled distance distribution with the lowest JS divergence from the experimental distribution was chosen for that model and spin pair. This approach was used to account for variability in the chiLife repacking method, which can introduce heterogeneity in modeled distributions due to the Markov Chain Monte Carlo (MCMC) process of adding label rotamers. A 3D array was created in the shape *models x spin pairs x distance distributions* for each non-negative matrix factorization (NMF) component (components and the usage of NMF is described in Wingler et al.^26^) modelling run. This was reshaped to yield a 2D array of dimensions *models x spin pairs * distance distributions*, such that each distribution in a single model is a part of the same vector. Each model in the 2D array was then ranked by the JS divergence or inverse dot product from the experimental data reshaped into a 1D array of connected distance distributions, where the top *n* models were summed to represent the experimental data.

Within the top *n* models, the secondary structure local to the label sites were analyzed qualitatively for alterations that indicate overfitting. In the presence of overfitting, the model selection criterion was adapted to include a filter for excluding the change in secondary structure. For the β2AR, approximately 600 of the models were over fit by showing a helix-to-loop transition at the cytoplasmic surface of TM6 at the site of the applied bias. These models were filtered out using HELANAL^74^ in MDAnalysis^75,76^ to calculate a local helix height sum of < 11Å from residues 267-275, leaving approximately 1200 models with the cytoplasmic end of TM6 in an alpha helix. No filter was used for the AT1R models. Following the filtering, the models were re-ranked according to the JS divergence or inverse dot product as described in the previous paragraph.

### DEER data used for model restraint generation

DEER data for β2AR 148/266 IAP were published in Lerch, et. al. 2020^18^. In this publication, the active conformation of TM6 populated by epinephrine + NB80 was represented as a gaussian centered at 50 Å, which was used herein as input data for modeling. DEER data for the AT1R was published in Wingler et. al 2019^26^, comprised of ten spin pairs and ten different liganded and unliganded conditions. NMF resolved four unique components (i.e., conformations) represented in the distance distribution data. Distance restraints from each of the four components were used here in modeling.

## Results

### Brief method overview

ProGuide presented here enables modeling of large global conformational rearrangements using multiple experimental distance distributions while minimizing structural overfitting errors. To achieve this, a three-block approach is taken: update block, biased MD block, and analysis block (**Figure 2**). In the update block (**Figure 2A**) the prior output model is used as an input, and an update to the target distance between the Cα atoms of the spin labeled residues is calculated based on the space within the experimental data not explored by the prior model. The updated Cα-Cα distance targets serve as restraints for the next modelling iteration. Using the biased MD block (**Figure 2B**), the distance update is driven by a learned linear bias potential applied globally and not restricted to the label sites. When incorporating datasets encompassing multiple spin pairs, the bias potential as a function of each pair is optimized simultaneously to allow for coordinated conformational rearrangements. After all update and MD blocks have completed, the models generated at each iteration of the bias-application block are binned, ranked, and a representative subset is selected in the analysis block (**Figure 2C**) for further structural and statistical analysis. Details of the method and implementation can be found in the *Methods* section.

### Activation of the β2AR

We demonstrate the ability of ProGuide to capture a large, complex conformational rearrangement using a single spin pair on the β_2_-adrenergic receptor (β2AR), which is a well-studied class A GPCR with available structures in various conformational states. Its structural rearrangements span up to 10Å Cα displacement in non-loop regions where maintaining secondary structure is crucial due to the prevalent allosteric communication between helices of GPCRs. Because β2AR has been extensively characterized, it serves as an excellent test system for this method. We used previously published DEER data on β2AR 148/266 IAP bound to agonist isoproterenol and nanobody-80 (Nb80)^18^ – an active state stabilizing nanobody – as the target distribution to model while beginning from an inactive β2AR conformation. This construct has been used in multiple DEER studies to report the conformational equilibrium of transmembrane helix (TM) 6, which exhibits the largest conformational rearrangement between inactive and active states.

Model generation of the β2AR was performed in six individual runs with 30 iterations each (180 total) using an inactive structure as the starting model. Following training and convergence of the distance update in the bias-application block, a production phase of 5 ns was used to allow relaxation around the bias restraints following the structural perturbation, totaling 900 ns of production simulation time. To assess how well the generated models matched the imposed distance restraints for each simulation, we compared the target Cα-Cα distances with the average Cα-Cα distances during the production phase. The RMSE between these values is 1.676 Å, indicating good agreement between the generated models and the target distances. A plot of the average production phase Cα-Cα distance vs the target Cα-Cα distance for each of the 180 iterations is shown in **Figure 3A**. Interestingly, the production phase distances are systematically lower than the target distances, likely due to the modeled distance change – an ∼10 Å increase in the Cα-Cα distance from the starting model. After assessing target distance convergence, we compared the simulated distribution of each model across iterations to the experimental distance distribution by JS divergence. This resulted in continuous improvements of the JS divergence (lower is better) from iterations 0 to 8 improving from >0.7 to ∼0.3 (Figure S2A). For the remainder of the modeling runs, the models explored JS divergences within the 0.1-0.6 range.

**Figure 3:**
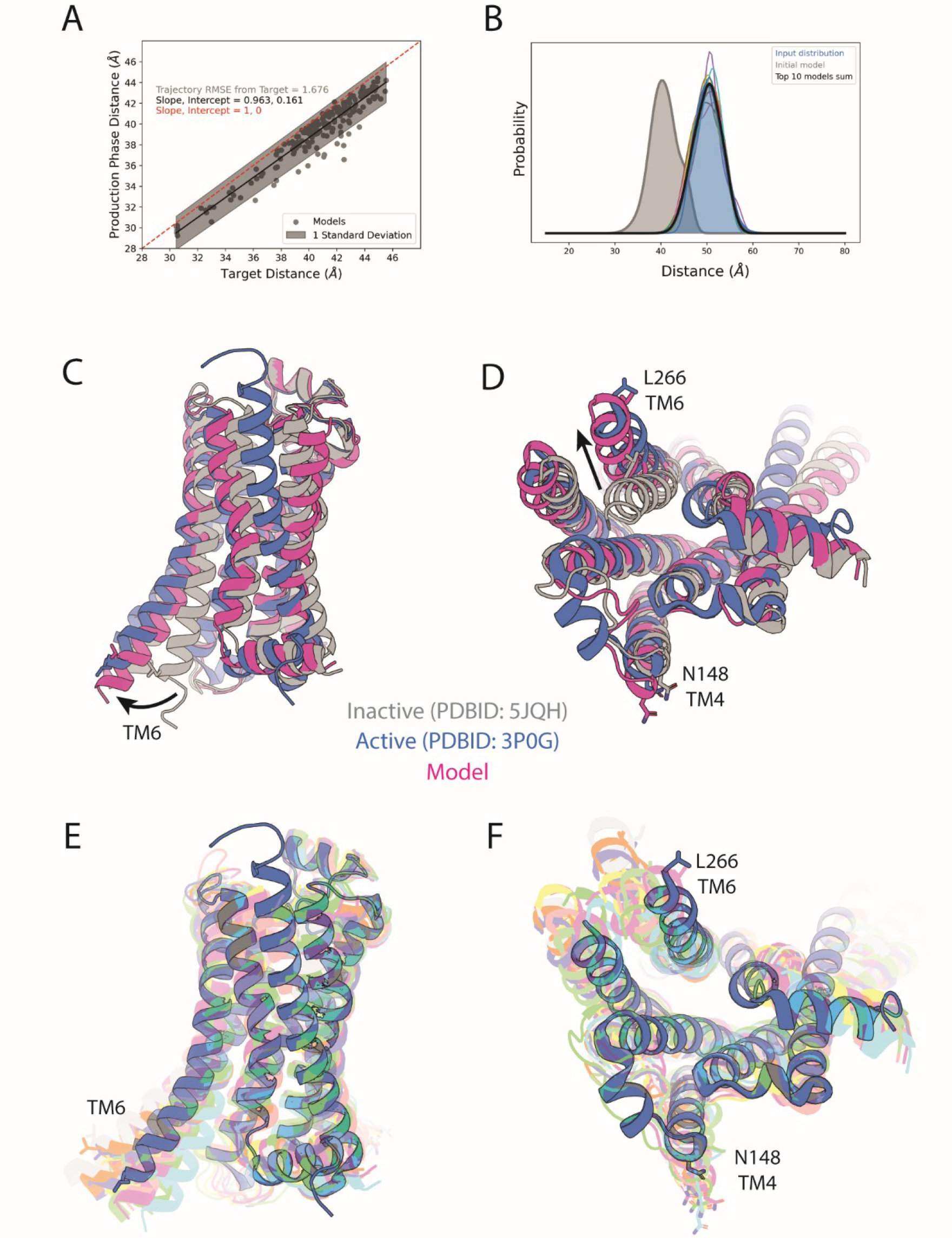
Modelling the outward active conformation of TM6 in the β2AR using a single IAP spin label pair at 148/266. A) Target Cα distance of the biased MD simulations vs. the Cα distance of the trajectory average of each modelling iteration. Data points (grey dots) are fit to a line (black) with one standard deviation in the fitting parameters (grey shade). A perfectly matched correlation between the trajectory average and the target distance would yield a line with slope 1 and intercept 0, modelled as a red line. B) Graph of the simulated chiLife distribution of the initial model (grey), the experimental data (blue), the simulated distributions of the top 10 models (multi-colored lines), and the sum of the top 10 models (black line). All distributions are area normalized. C-F) Cartoon representations of the starting model (grey), active crystal structure of the β2AR (PDB ID: 3P0G), a representative model (magenta) from the top 10 models, and the top ten models shown as transparent cartoons (multi-colored). Residues N148 (TM4) and L266 are shown as sticks. Intracellular loop (ICL) 3 has been truncated in all models shown here for visual purposes.

Ten frames from the last ns of the production phase (one every 0.1 ns) from each iteration were used as models in the analysis block, for a total of 1800. After simulating the rotamer distributions using chiLife, the models were ranked by JS divergence and a final ensemble comprised of the top ten models was used for structural analysis. Simulated distance distributions for each of the top ten models demonstrate good agreement with the target distance distribution, and the linear combination of the top ten models achieves a JS divergence of 0.04, indicating nearly perfect overlap with the experimental data (**Figure 3B**). A representative model is overlayed with β2AR inactive state (i.e., starting model) and active state (i.e., target structure) in **Figure 3C and 3D**, respectively, highlighting the ∼10 Å shift of TM6 from an inward inactive position to an outward active position while minimizing local secondary structure changes. Though all top 10 models show similar outward rearrangements of TM6, some models deviate further from the overlayed β2AR crystal structure demonstrating conformational heterogeneity that is still consistent with the experimental data (**Figure 3E/F**). To quantify this heterogeneity, a Cα RMSF plot was created using the top 10 models (**Figure S2B**), where the residue specific Cα deviation from an average model is easily visible. In TM6 from residues 261-280, the mean Cα RMSF is 2.35Å and the RMSF at site 266 is 2.13Å.

### Biased conformational states of the AT1R

To showcase ProGuide’s capability to model conformational rearrangements using restraints from multiple spin pairs, we apply it to the AT1R utilizing DEER data reported by Wingler et al.^26^ The DEER dataset comprised ten spin pairs distributed across the cytoplasmic surface, which were used to monitor conformational changes induced by ligands with distinct efficacy profiles (i.e., inverse agonism, balanced agonism, β-arrestin-biased agonism, or Gq-biased agonism) to provide insight into the structural basis for biased agonism. The prior study identified a set of four distinct conformations via DEER data deconvolution into four non-negative matrix factorization (NMF) components, whose relative populations are modulated by ligands in correlation with their functional profiles. To visualize and compare the different conformations, spin density clouds were calculated and projected onto published structures without generating atomistic models from the DEER data. Here, we used distance restraints from the AT1R DEER data to generate complete structural models of each of the four conformations. Using an inactive state starting model, eight modelling runs with a maximum of 30 iterations were performed for each component. Because there is the potential for an individual run to get stuck in a local energy minimum, we used multiple replicates to increases how much of the conformational space available to the receptor was explored. The top 50 models were combined with equal weights and used to represent the final distributions for each component (Figure 4A).

**Figure 4:**
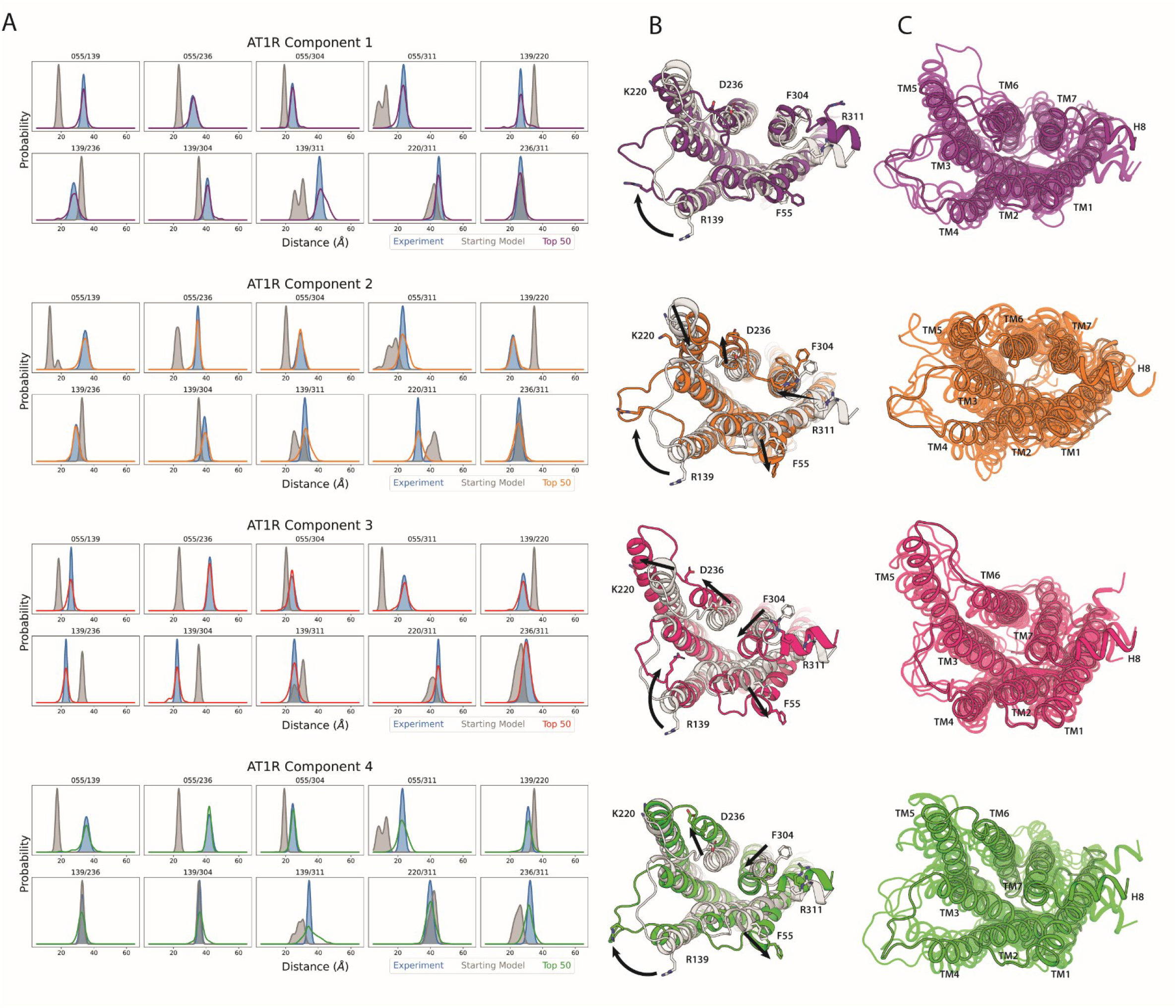
Results from modelling NMF components of the AT1R from the data in Wingler et al. 2019. A) Graphs depicting the NMF conformation/input data (blue) vs. the sum of the simulated spin label distributions of the top 50 models (colored).Starting model simulated distributions are represented in grey. B) Representative AT1R conformational models (color) overlayed with the starting model generated from an unbiased MD equilibration of the antagonist bound AT1R structure (PDB ID: 4YAY). Each model was selected from the top 5 models ranked by their closeness to the NMF component data. Label sites are shown as their native residues represented by sticks. Arrows indicate major structural changes observed in the overlayed structures. C) Ensemble of the top 5 models in each NMF conformation. Models highlighted with a black silhouette are the models representing the conformational components in panel B.

Excellent agreement is observed between the most probable distance for the modeled and experimental distributions (Figure 4A), even in cases where this required a shift of ∼20 Å from the starting model (e.g. 55/139-V1 in component 3). This highlights the ability of ProGuide to produce coordinated structural rearrangements driven by multiple distance restraints. Good agreement is also found in the widths of most modeled and experimental distributions (Figure 4A), with the exception of the 139/311-V1 pair. Residues 139 and 311 are found in ICL2 and helix 8, respectively, which are highly dynamic regions of the receptor. A comparison of the target distances with average production distance between labeled Cα residues in the AT1R conformations 1-4 showed RMSE values of 0.993 Å, 1.398 Å, 0.863 Å, and 1.379 Å, respectively (Figure S5).

Overlays of the starting AT1R model with representative models for each conformation are shown in Figure 4B to illustrate the structural rearrangements on the cytoplasmic surface. Major structural changes for each conformation are highlighted (black arrows, Figure 4B) by the Cα atom of the spin site in the corresponding loop or helix by changes >3 Å. All distances referenced below indicate changes in spin site Cα positions between the starting model and the aligned representative model conformation. The primary and most notable rearrangement is found in intracellular loop 2 (ICL2), driven by an increase in the distance for the 55/139-V1 spin pair. Conformations 1, 2, and 4 exhibit an outstretched ICL2 that is displaced from the cytoplasmic bundle, whereas ICL2 is positioned near the helical core in conformation 3. Conformation 1 shows a 3.2 Å outward shift in residue 304 (TM7) as the primary conformational rearrangement. Conformation 2 displays a 3.6 Å outward shift in residue 55 (TM1), a 3.9 Å inward displacement of residue 220 (TM5) towards the center of the helical bundle, a 6.9Å outward and upward shift of residue 236 (TM6), a 3.7 Å outward displacement of residue 304 (TM7), and a 13.2Å shift in residue 311 (H8). Interestingly, conformation 2 is the only conformation that showed a consistent displacement of residue 311 (H8) away from the plane of the membrane (Figure 4B, Figure S8), as well as the only conformation to show an upward displacement in TM6 (Figure S8). Conformation 3 shows a significant outward displacement of 7.7 Å in residue 236 (TM6), 4.9 Å in residue 220 (TM5), and 7.9 Å in residue 55 (TM1). Conformation 3 models also show a significant inward transition of residue 304 (TM7) by 7.1 Å, likely accommodated by the outward shift in TM1. The starting model 55/304 Cα-Cα distance lies within the range of the conformation 3 distance range for this spin pair (Figure S3), yet the overlayed models show a significant change in the two helices containing these residues due to the concomitant changes in other spin pairs involving residue 55 (TM1) and 304 (TM7). Conformation 4 shows a significant outward displacement of 6.9 Å in residue 236 (TM6) and 6.4 Å in residue 55 (TM1), whereas residue 304 (TM7) shows a 5.3 Å shift towards the helical core, again likely accommodated by the outward movement in TM1. To aid in quantifying the heterogeneity at these sites, the RMSF of each label site Cα for the top 50 models of each component are listed in **Table S3** where the conformational rearrangements from the starting model mentioned above (measured from the starting state) are all greater than their respective RMSF values at each site (RMSF plot in Figure S4), highlighting that these rearrangements are significantly different than the starting state. The top 5 models generated from each component (Figure 4C) demonstrate the heterogeneity present in the ensemble for each conformation.

Principal component analysis (PCA) conducted on the coordinates of the protein backbone for the top 50 models generated from each component (Figure 5A) visualizes the non-random pattern of model dispersion – in PCA space – from the same starting model. The results demonstrate that the models undergo *global* conformational rearrangements from the linear bias derived from distance restraints on a small subset of atoms within the protein system. When projected on to the protein backbone, PC1 involves most of the cytoplasmic surface of the receptor with inward and outward movements of ICL2, TM1, TM5, TM6, TM7 with respect to the center of the helical bundle, as well as an up and down motion of H8 with respect to the membrane, all of which were mentioned in major rearrangements in the above paragraph. PC2 involves more subtle motions of the cytoplasmic helices; primarily lateral motions in H8 coupled with minor motions of TM6. PCA conducted on the Cα atom coordinates of the individual label sites exhibit a similar pattern to the backbone PCA. As expected, the label site Cα PCA separates into discernable clusters (Figure 5B), due to bias placed on the system as a function of the distance between label site Cα atoms. The cluster representation similarity between the backbone PCA and label site PCA demonstrates that while the label sites are specifically targeted to meet an experimental restraint, the rest of the system responds to the bias in a coordinated manner.

**Figure 5:**
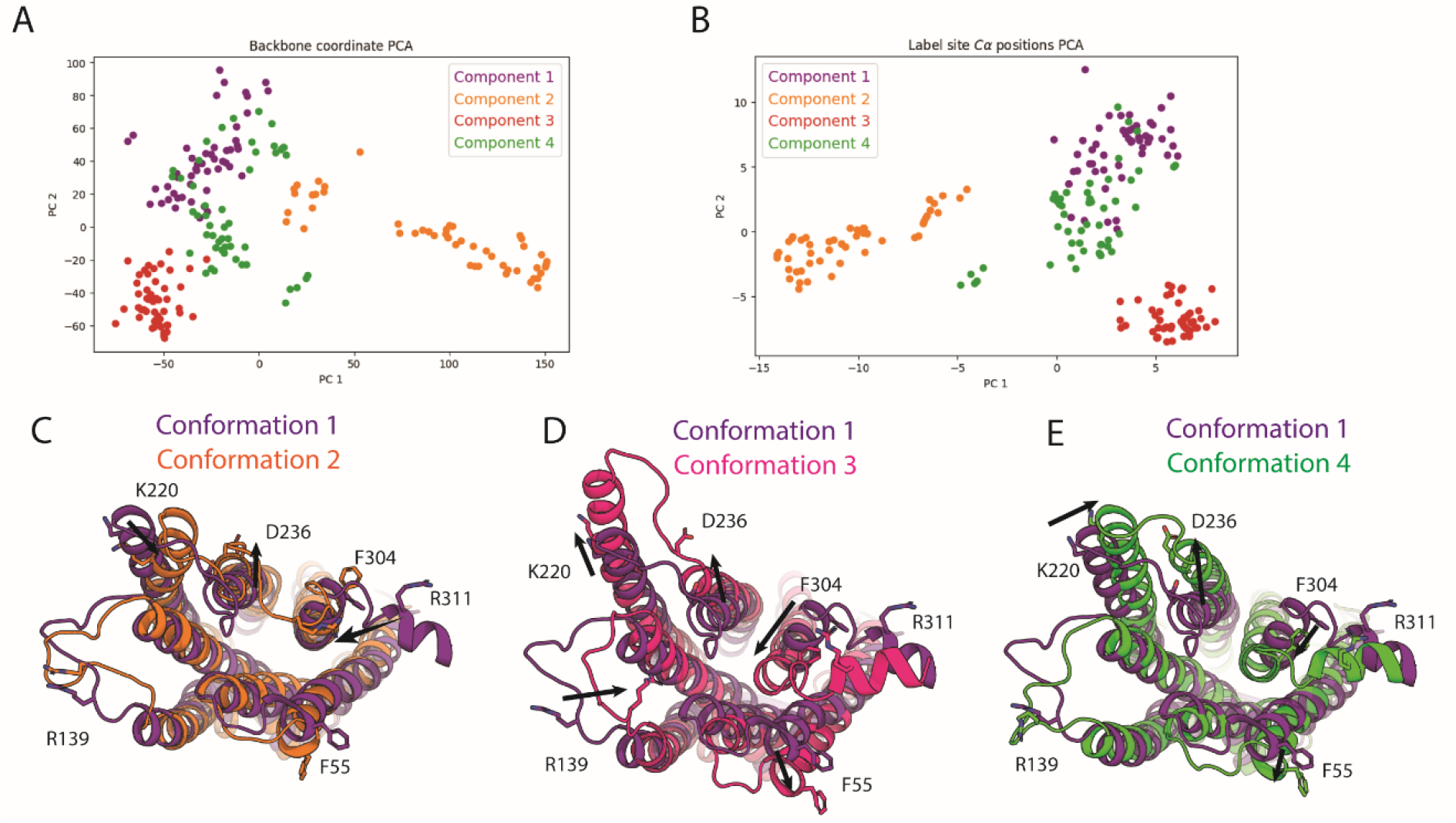
Model and motif analysis of the AT1R models generated from NMF components. A) PCA of AT1R model backbone atom coordinates and B) AT1R model label site Cα atom coordinates (lower). C-E) Comparison of the conformation 1 model to conformation 2 (C), conformation 3 (D), and conformation 4 (E). Models are shown as cartoons, native label site residues are shown as sticks, and relevant conformational rearrangements are highlighted with a black arrow.

Overlays of representative models for conformations 2-4 with a representative model of conformation 1 are shown in Figure 5C-E to highlight the structural differences between conformations. To highlight relevant changes between the conformation models here, the overlayed representative model Cα change must be >3Å compared to conformation 1 for the Cα change of interest. Conformation 2 shows a 4.9Å inward shift of TM5, a 6.9Å outward and upward shift of TM6, and a 17.7Å displacement of H8 compared to conformation 1 (Figure 5C). Conformation 3 shows the most differences from conformation 1 with outward movements of TM1, TM5, and TM6 by 6.9Å, 3.2Å, and 6.5Å, respectively (Figure 5D). In comparison to conformation 1, the conformation 3 model also shows an inward shift of 9.6Å in residue 304 (TM7) and a 9.7Å inward positioning of residue 139 (ICL2) (Figure 5D). Lastly, conformation 4 shows a 3.9Å shift of residue 220 (TM5) approaching TM6, while residue 236 (TM6) moves 6.3Å into an outward position (Figure 5E). Residue 304 (TM7) adopts an inward position, moving 7.8Å towards the helical bundle and residue 55 (TM1) accommodates for this by shifting 5.3Å away from the helical core.

To demonstrate motif-specific conformational rearrangements, we highlight the relatively large movements of the NPxxY motif (N7.49 to Y7.53, Ballesteros-Weinstein numbering) and more subtle residue rearrangements of the DRY motif (D3.49 to Y3.51), both of which are conserved among class A GPCRs. Figure 6A shows Cα distances between P299 (P7.50) and L70 (L2.46) of representative models for conformation 2 and conformation 3; a decrease in this distance indicates an inward displacement of TM7 associated with GPCR activation^77,78^. The L70-P299 Cα distance was measured across all models and plotted (Figure 6B); statistically significant differences were observed between each of the different conformational ensembles (p-value < 0.05, Table S2). For conformation 3, 68% of the top 50 models fall below the 7Å threshold for β arrestin bias positioning^78^ (black dotted line, Figure 6B).

**Figure 6:**
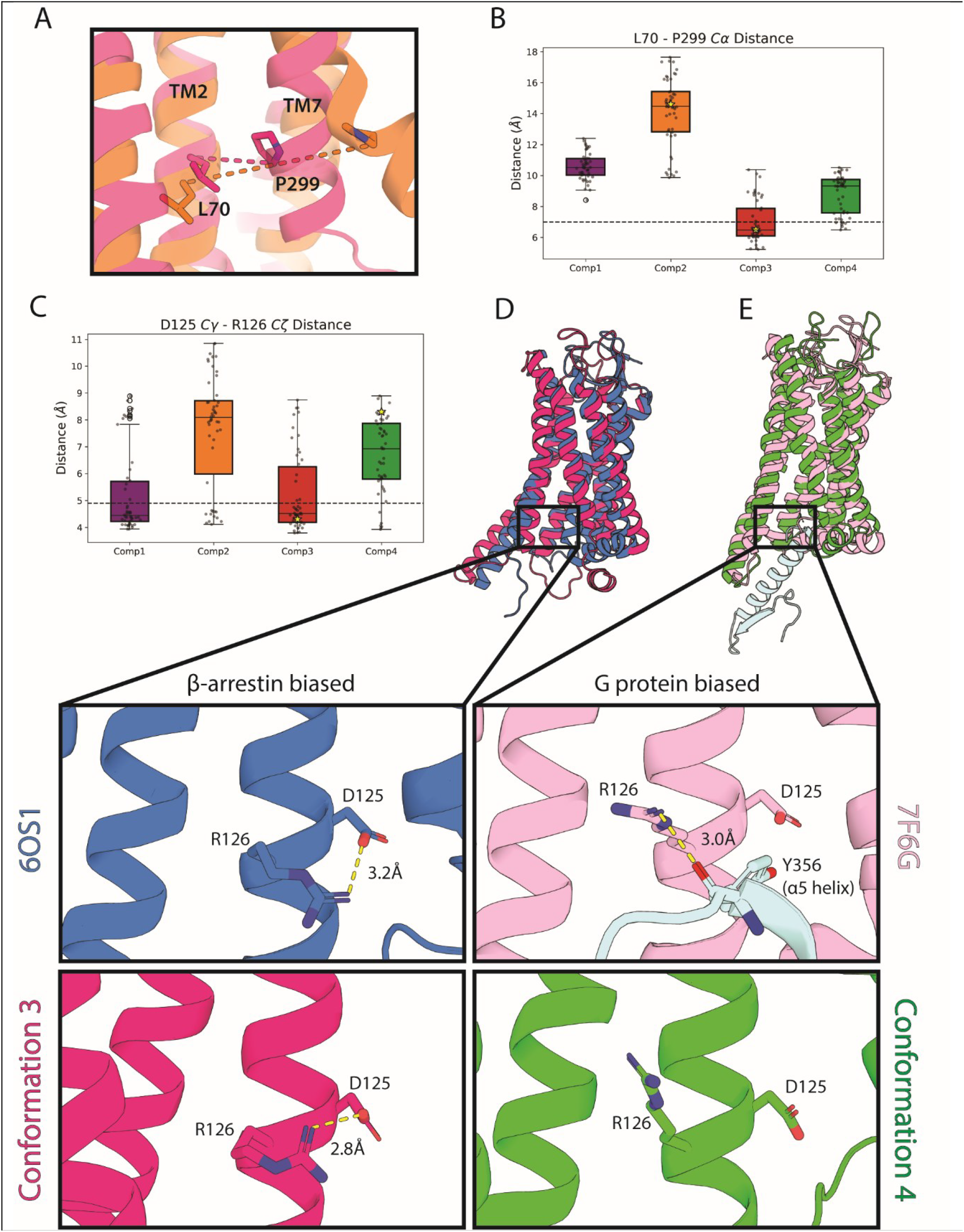
Models show NPxxY and DRY motif changes consistent with published X-ray crystallography and cryo-EM structures. A) L70-P299 Cα atom distance as a monitor of NPxxY movement of the AT1R models. Representative models of conformations 2 (orange) and 3 (green) are shown as cartoons, and the distance between the Cα atoms are highlighted by dashed lines. B) Box and whisker plot for L70-P299 Cα atom distance of models generated in each conformation. The median value is indicated by the center line and lower and upper bounds indicate the first and third quartile values. Upper and lower bounds of the whiskers indicate the furthest datapoint lying within -1.5 IQR and +1.5 IQR, respectively. Black dots represent data points, open circles represent outliers, and the stars are the values of the models in panel A. The black dashed line at 7Å represents the upper limited considered for β-arrestin-biased conformations of the NPXXY measured by L70-P299 distance.^78^ C) Box and whisker plot of the distance between D125 Cγ and R126 Cζ of the DRY motif for the models in each conformation. The median value is indicated by the center line and lower and upper bounds indicate the first and third quartile values. Upper and lower bounds of the whiskers indicate the furthest datapoint lying within -1.5 IQR and +1.5 IQR, respectively. Black dots represent data points, open circles represent outliers, and the stars are the values of the models in panels D and E. The black dotted line is the distance between D125 Cγ and R126 Cζ atoms of 6OS1. D) Overlay of the TRV023 (β-arrestin-biased ligand) bound AT1R X-ray crystal structure (PDBID: 6OS1) with a representative model of conformation 3. Blow up panels show the DRY motif of each structure/model. A salt bridge interaction between D125 and R126 is shown by the yellow dashed line. E) Overlay of the AT1R-Gq complex bound to Sar1-AngII (PDBID: 7F6G) with a representative model of conformation 4. The α5 helix of the Gq protein is shown in pale cyan. The upper panel shows R126 of the AT1R interacting with Y356 of the Gq α5 helix, shown by a yellow dashed line.

Rearrangements in the residues of the DRY motif were monitored using the distance between D125 Cγ and R126 Cζ atoms (Figure 6C). Statistically significant differences are observed across all conformations (p<0.05, Table S4), except for conformations 1 and 3. A distance below 4.9Å results in D125 oxygen and R126 nitrogen atoms within range to form electrostatic interactions or a salt bridge, which we refer to as a “closed configuration” of D125-R126 (black dotted line, Figure 6C). The closed configuration is observed for the β-arrestin-biased conformation 3 model, consistent with the crystal structure of AT1R bound to the β-arrestin-biased ligand TRV026 (PDB: 6OS1, Figure 6D). An “open configuration” of D125-R126 is observed for the Gαq-biased component 4 model, in good agreement with a cryo-EM structure of the AT1R-Gq complex^79^ (PDB: 7F6G, Figure 6D). Among the top 50 models generated for each component, 66% of component 1 and 3 models were in a closed D125-R126 configuration, whereas an open configuration was observed for 76% and 88% of component 2 and 4 models, respectively (Figure 6C).

## Discussion

### ProGuide performance

Here we describe ProGuide, a framework for modeling global structural rearrangements in proteins using DEER-derived distance restraints from multiple spin pairs. In iterative biased molecular dynamics simulations, incremental changes are driven in the distances between Cα residues based on a comparison of experimental DEER distributions with simulated distributions generated via *in-silico* spin labeling with chiLife. The resulting ensemble of models is then subjected to a selection process to select the models that best recapitulate the DEER data. We illustrate the power of this method using published DEER data on a pair of GPCRs. First, a ∼10 Å outward rigid-body motion of TM6 is modeled in the β2AR using DEER data from a single spin pair. In a second application, the power of ProGuide is illustrated using DEER data of multiple spin pairs from a study of biased agonism in the AT1R. The AT1R models generated here display tertiary rearrangements from the starting model as well as residue-level microswitch rearrangements. Below, we discuss the modeling accuracy in terms of model fidelity to the assigned target distance and simulated distribution fit to the experimental data.

Each iteration of model generation is guided by a trained linear bias potential to a target distance and then maintained through a 10 ns production simulation that allows the entire system to relax around the bias potential. Excellent agreement is observed between target and observed Cα distances in production simulations for all modelling runs performed here (RMSE = ∼0.9-1.7 Å; Figures 2A and S5). Importantly, these values demonstrate ProGuide’s ability to induce and preserve structural rearrangements throughout the modeling process, achieving sub-angstrom error in some cases. The ability to resolve subtle structural rearrangements with high confidence enables the use of ProGuide to reveal small but functionally significant conformational changes.

The local secondary structure at each label site is maintained for >60% of models in the β2AR ensemble and for all models in the AT1R ensemble. This is crucial to avoid overfitting the experimental data by denaturing local secondary structure, which may produce physically unreasonable models. The simulated distributions for the unrefined β2AR and AT1R ensembles displayed excellent coverage of the experimental distributions, demonstrating the ability of ProGuide to explore the entire restraint space. Refinement of the ensemble by JS distance improves the fit and selects for models that best agree with the experimental data (Figure S6).

The excellent agreement between simulated and experimental distance distributions achieved through physically reasonable structural rearrangements in the protein structure are due largely to placing the bias on the backbone of the protein, rather than the spin labels. As a result, optimization of the nitroxide distributions is achieved by movements in the underlying model protein, rather than distortion of spin label side chain rotamers. Additionally, rearrangements of the protein are optimized through physics-based MD, capturing allosteric conformational rearrangements not explicitly reflected in the experimental restraints and producing more accurate conformations.

ProGuide presents broad customizability that facilitates application to a wide range of biological targets. The framework is built on the popular GROMACS MD simulation software, enabling tailoring of the simulation environment (e.g., soluble protein, membrane protein, lipid composition, pH) using standard tools. Ligands or interacting proteins can also be added to the simulations during the modelling process. Multi-protein systems with spin label pairs across different protein chains or on a singular chain can be accommodated with no changes in the underlying framework. The adaptability and flexibility of chiLife opens the doors to vast possibilities in modeling proteins using experimental DEER data as restraints. Currently, chiLife hosts MTSSL/R1, R7, V1, IAP, TRT, and others, with the option to use labeling methods implemented in MMM^80^ or MTSSLWizard^81^, along with the chiLife methods.^50^ New labels can be added in a straightforward manner, and bond dihedrals can be customized to generate custom rotamer libraries for existing or new spin labels. This functionality enables modeling of novel spin labels that are developed and used in DEER experiments but have not yet been parameterized for MD simulations.

The distance restraints used here are derived from probability distributions of distances obtained by fitting DEER dipolar evolution data. In the case of the AT1R, DEER data collected for ten spin pairs under different ligand conditions revealed significant heterogeneity in the form of multi-modal distance distributions observed in many cases. Correlated ligand-dependent changes across the different spin pairs were identified via NMF of the distance distributions, and the resulting components were used in this work as experimental restraints to generate structural models of distinct conformations represented in the DEER data. While not trivial, it is more tractable to identify correlated features across spin pairs using distance distributions rather than dipolar evolution traces. As such, it is preferable to use distance distributions as experimental restraints to model distinct conformations represented in datasets with multiple spin pairs. Additionally, the use of distance distributions provides flexibility to exclude distance peaks arising from off-target labeling, oligomerization, regularization routines, or other experimental imperfections.

### Generated models of AT1R capture biased conformations

Wingler et. al 2019^26^ identifies the presence of four distinct conformations of the AT1R through NMF of ten DEER spin label pairs in ten ligand conditions. NMF deconvolutes data into a user-identified number of components and their contributions to each piece of data (i.e., each ligand condition from the DEER data). Four NMF components were identified in the DEER data that translated to four separate conformations of the AT1R. From these identified components, nitroxide spin positions were modelled in 3D space and aligned to simulation snapshots of the AT1R with modelled V1 labels. Here, ProGuide yields atomistic protein models corresponding to four conformations from each of the 4 NMF components described in Wingler et. al 2019^26^. We focus on correlations between conformational changes at the intracellular surface with functional selectivity of the ligands used in the original study.

#### Conformation 1

Of the 4 conformations, conformation 1 is most similar to the starting model, which was generated via an unbiased MD equilibration of the crystal structure of AT1R bound to a precursor of the inverse agonist candesartan (ZD7155, PDB: 4YAY). Unsurprisingly, conformation 1 is most highly populated in the DEER data collected on receptor bound to inverse agonists candesartan and olmesartan. Both the starting model and conformation 1 exhibit the structural features of a canonical inactive conformation, including an inward position of TM6. Despite the overall structural similarity, there are major differences in their distance distributions, with 5 of the 10 spin pairs exhibiting >10 Å difference in the most probable distance (Fig. 4A). This discrepancy illustrates that significant changes in distance distributions can result from relatively minor structural rearrangements of the protein. To achieve the excellent overlap observed in the distance distributions for conformation 1 and component 1, the primary structural rearrangement from the starting model to conformation 1 is an unraveling of the cytoplasmic end of TM4 and a repositioning of ICL2 (Fig. 4B); only small rigid-body movements are observed in the transmembrane helices.

#### Conformations 3 and 4

Several AT1R structures were published after the AT1R DEER data analyzed here were originally reported, including a cryo-EM structure of AT1R bound to Gq (PDB ID: 7F6G) and crystal structures of AT1R bound to β-arrestin-biased agonists TRV023 (PDB ID: 6OS1) and TRV026 (PDB ID: 6OS2) with active state-stabilizing nanobody AT110i1. Comparison of conformation 3 and 4 models with these structures^79,82^ reveals deeper, residue-level information obtained using ProGuide.

An overlay of a representative model of conformation 4 with the AT1R-Gq complex structure (Figure 7A) demonstrates a remarkably similar structure at the cytoplasmic surface of the receptor. The AT1R-Gq complex exhibits an open DRY configuration, allowing AT1R R126 to stabilize the Gq α5 helix of by interacting with the carbonyl oxygen of the Y356 backbone (Figure 6E)^79^. Similarly, an open DRY configuration is found in conformation 4 models, positioning R126 for interaction with the α5 helix. Conversely, the crystal structure of AT1R bound to TRV023 and nanobody AT110i1 shows a closed DRY configuration that is hypothesized to clash with Gq α5 helix engagement to the AT1R^82^. Mutation of R126 to alanine reduces the relative Gαq activation by AT1R without significantly affecting β arrestin activation^83^. Conformation 3, which is associated with β-arrestin bias, also shows a closed DRY configuration (Figure 6D). With this information, we propose that the closed configuration of R126 (Figure 6D) promotes β arrestin selectivity by inhibiting Gq binding.

**Figure 7:**
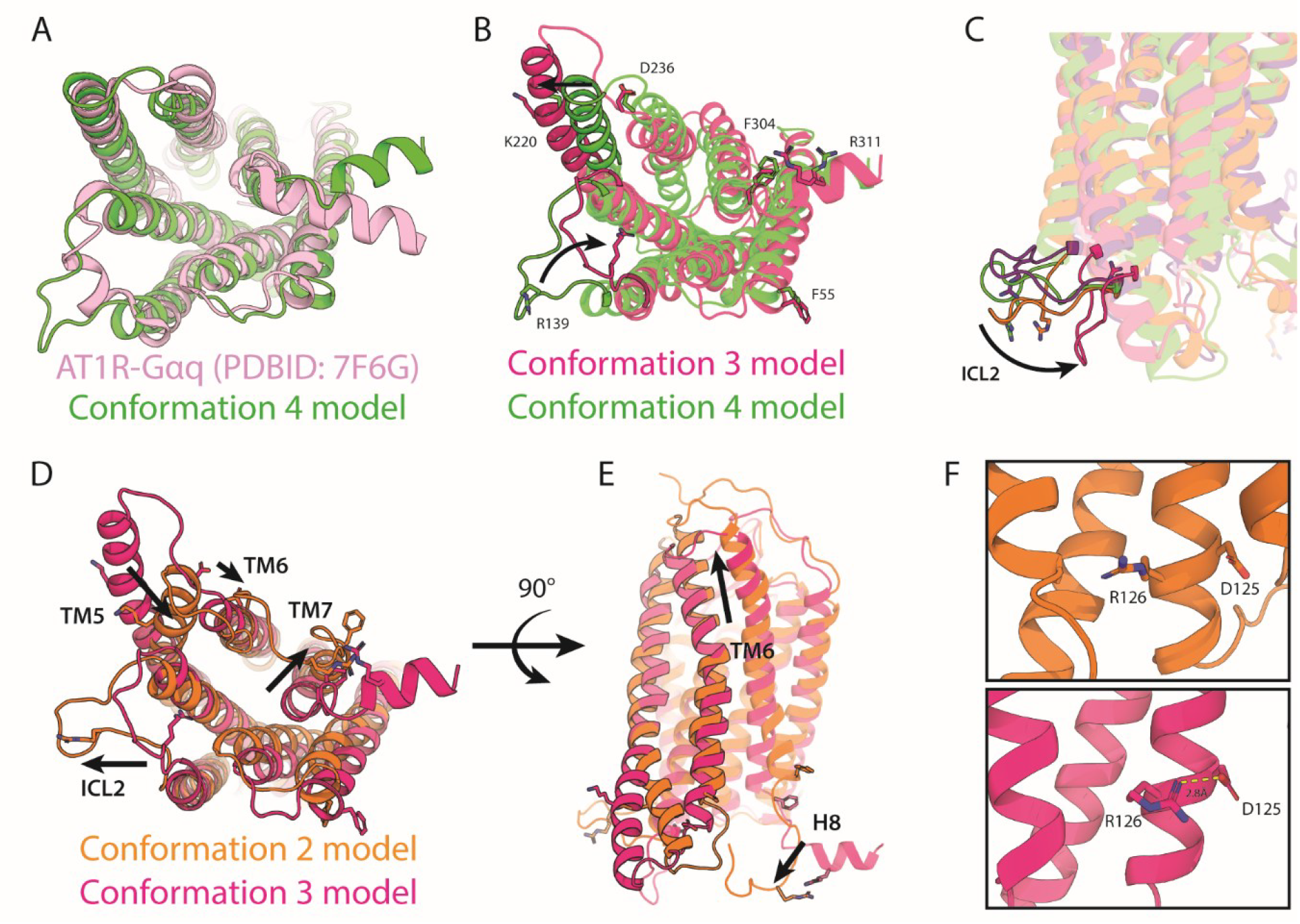
Comparison of highly relevant conformational rearrangements between models and comparison with previous cryo-EM structures. A) Comparison from the cytoplasmic surface of Gq biased conformation 4 with the cryo-EM structure AT1R-Gαq complex (PDB ID: 7F6G, Gαq removed for clarity). B) Comparison from the cytoplasmic surface of conformational rearrangements (black arrow) of β-arrestin biased conformation 3 (magenta) and Gq biased conformation 4 (green). C) A view from the membrane plane showing the inward conformational rearrangement in ICL2 of conformation 3 compared to the other conformational models. D) Comparison of β arrestin biased models derived from conformation 2 (orange) and conformation 3 (magenta) from the cytoplasmic surface. Helical rearrangements from conformation 3 model to the conformation 2 model are shown by arrows. E) Comparison of β arrestin biased models derived from conformation 2 (orange) and conformation 3 (magenta) from the membrane plane. Helical rearrangements from conformation 3 model to the conformation 2 model are shown by arrows. F) DRY configurations showing an open configuration for the conformation 2 model (orange, top) and a closed configuration for the conformation 3 model (magenta, bottom). The distance between the R126 nitrogen atom and D125 oxygen atom is highlighted by a yellow dashed line.

Overlaying representative models of the β-arrestin-biased conformation 3 with that of Gq-biased conformation 4 highlights the primary structural differences to be an inward conformation of ICL2 and an outward conformation of TM5 in conformation 3 (**Figure 7B**). These differences were discussed in Wingler et. al 2019^26^ as well, though a subtle but statistically significant rearrangement in TM7 was missed. While both conformation 3 and 4 models show inward movement of TM7 indicative of GPCR activation, only conformation 3 meets the sub-7 Å L70-P299 distance threshold (dashed line, Fig. 6B) identified for β-arrestin-bias in a previous computational study^78^ of AT1R bound to each of the ligands used in the Wingler et al. 2019^26^ study. We postulate that – in addition to rearrangements in ICL2 and TM5 – the position of TM7 is a key indicator for G protein vs. β arrestin bias, which is not immediately apparent from inspection of DEER data or rotamer cloud modelling alone.

Taken together, we can correlate the DRY configuration, NPxxY position, and other helical/loop movements at the cytoplasmic surface in conformations 3 and 4 to functional selectivity. The representative models for conformation 4 indicate selectivity for Gq likely arises from positioning ICL2 and TM6 outward to not obstruct the cytoplasmic binding surface, TM7/NPxxY adopting an inward position, and the DRY adopting an open configuration to stabilize the α5 helix as Gq interacts with the AT1R. Conformation 3 models show that Gq binding may be interrupted and bias towards β arrestin generated by the inward movement of ICL2, TM7/NPxxY moving to a fully inward conformation, and DRY adopting a closed configuration that clashes with α5 helix stabilization. These results agree with those from previous alanine scanning studies^83^ showing that mutations near/in ICL2 and in the DRY motif displayed preferential engagement of β-arrestin over Gq.

#### Conformation 2

A second β-arrestin-biased conformation was described in Wingler et. al 2019^26^ based on NMF component 2, used here to generate conformation 2 models. Compared to conformation 3, conformation 2 shows an outward ICL2 position, inward TM5 position, slight inward movement of TM6, open configuration of the DRY motif, and outward position of TM7/NPxxY (Figure 7D). In conformation 2, TM6 also shows a vertical displacement and H8 is vertically displaced from the membrane (Figure 7E, Fig S8). Conformation 2 thus represents an alternate β-arrestin-biased conformation; one that has not been previously reported. Some of the unique structural features of conformation 2 can be inferred from DEER data. However, subtle details of the helical arrangements, vertical displacement of TM6, displacement of H8 from the plane of the membrane, and the addition of the DRY configuration provides crucial structural details into this alternate β-arrestin-biased conformation that could not be postulated from DEER data alone.

Interestingly, the crystal structure of AT1R bound to TRV026 (6OS2) is identical in helical arrangement to the crystal structure of AT1R bound to TRV023 (6OS1) (Fig S7, RMSD: 0.524), which is inconsistent with NMF analysis of the DEER data and the models reported here. Component 2 was most highly populated (NMF amplitude factor) in AT1R bound to TRV026, and component 3 was the dominant component for AT1R bound to TRV023. As described above, conformations 2 and 3 are structurally distinct, yet the crystal structures of AT1R bound to TRV023 and TRV026 do not indicate distinct effects of these ligands at the cytoplasmic surface of the receptor. This may be due to the fact that each of these structures was solved for AT1R bound to the same active state-stabilizing nanobody,^82^ which has a significant impact on the final stabilized conformation present in the crystal. The results shown here illustrate that DEER spectroscopy coupled with ProGuide can reveal structural effects of ligand binding that are obscured by the use of nanobodies or other fiduciaries commonly used in cryo-EM and crystallography.

## Conclusions

Here, we have developed ProGuide, a framework to effectively model functionally relevant conformations of proteins using DEER-derived distance restraints. This framework presents a tremendous level of flexibility, enabling customization of the simulation environment to suit different proteins, labels, and solvation conditions. The IMRSE algorithm efficiently produces update target distances based on unexplored model space. The BRER engine implements the target distances and allows the for the tuning and efficient use of a linear bias potential to impart coordinated structural rearrangements on the entire protein system as a function of the experimental data. Importantly, the labels are not present when the bias potential is applied to the system, leaving the rotameric distributions free from distortion. This approach enables elucidation of protein conformations associated with specific conditions (e.g., binding of ligands with various efficacies), offering deeper insights into protein function. Using ProGuide with previously published DEER data, we report Gq- and β-arrestin-biased conformations of the AT1R, including a completely novel β-arrestin-biased conformation. Structural insights from these models include tertiary structural rearrangements as well as residue-level rearrangements, allowing for the analysis of crucial microswitch motifs as they relate to biased signaling. These results illustrate the potential for ProGuide to aid in structure-based drug discovery efforts of GPCRs and beyond.

## Acknowledgements

Research reported in this publication was supported by the National Institute of General Medical Sciences of the National Institutes of Health under award number R01GM135581 (M.T.L.) and R01GM138444 (P.M.K). This research was completed in part with computational resources and technical support provided by the Research Computing Center at the Medical College of Wisconsin. The content is solely the responsibility of the authors and does not necessarily represent the official views of the National Institutes of Health. We are grateful for Maxx Tessmer’s consultation in the use of chiLife.

