## Supplementary Information for "ProGuide: a flexible framework for modeling global conformational rearrangements in proteins using DEER-derived distance restraints"

Toy model system used for conceptualization and parameter development

A simplistic two-point model system was used to conceptualize the algorithmic workflow with incorporation of the momentum (ρ) and learning rate (k) parameters. The toy model system can be seen as two points on a line with each point representing a Cα atom. The distance between the two is analogous to the distance between Cα atoms that are seen in full molecular simulations. Because spin labels are used to measure distances in DEER, a distance factor $d_f$ is added to the distance between points Cα to represent this. Taking the information of the toy system so far, a mean distance between spin labels in this system is defined by

$$\overline{u} = C\alpha + d_f$$

*(S 1)*

The orientation between spin labels can be assumed to be approximately fixed for linear motions with no rotations involved, so the distance factor is arbitrary and is only used as an extension representing spin label length, showing that Cα distance metrics can be updated and inform the rest of the probability distribution from these calculations. The spin label probability distribution is assumed to be Gaussian at solvent exposed sites (most often used for DEER studies). As such, a Gaussian model with parametric extension is used where $\overline{u}$ is the center of the distribution and a width is defined by the standard deviation σ

$$P(r) = e^{\frac{-(r-\overline{u})^2}{2\sigma^2}}$$

*(S 2)*

This probability distribution expression for the spin labels in this toy model represents the role of chiLife in the full molecular workflow. After performing a distance update calculation towards some experimental distribution, the updated model distribution center is now defined as

$$\overline{u_{new}} = C\alpha + \Delta C\alpha + d_f$$

*(S 3)*

Where ΔCα is the Cα update as described in the Methods section and is added to Cα to represent a target Cα distance that is fed into the biased MD block. The assumption is made that the target distance from the biased MD block is met, so in this toy model system Cα+ΔCα is used as the initial Cα in the next update iteration and formal representations of the training, convergence, and production phases in the biased MD block are dropped. Despite a direction-guided update, corrections towards the experimental distribution will not be perfectly linear due to the stochastic nature of molecular systems. To account for such stochasticity, a stochastic factor $s_f$ is introduced at each update step, where $s_f$ is randomly drawn from a uniform distribution of values from -2 to 2. The probability distribution for the updated model, accounting for $s_f$, can now be expressed as

$$P_{new}(r) = e^{\frac{-(r-(\overline{u_{new}}+s_f))^2}{2\sigma^2}}$$

*(S 4)*

The updates and toy system modeling follows the same workflow as presented in the Methods, where a number of iterations is performed and the previous model Cα+ΔCα distance is used as the input model Cα.

The toy model system here was sufficient to explore parameters and methods that may lead to improved exploration of experimental distribution space. In this way, momentum and learning rate parameters were introduced successfully and ultimately incorporated into the presented workflow.

The learning rate was incorporated early in the development as a means to prevent abrupt, high-energy bias potentials that would result in structural overfitting by local denaturation. By taking a fraction of the difference between the experimental distribution and modeled distribution, an incremental and effective structural transition could be attained while allowing the model to equilibrate to the applied bias at each step. A learning rate of 1 is the case in which no increment is taken, but rather a predicted “jump” directly to the explored space is initiated. This case leads to instability as visualized in Figure S9A. An optimal learning rate will aid in exploration of the experimental ensemble. The effect of changing this parameter is also shown in Figure S9A.

The momentum parameter enables prior model distributions to be accounted for during the distance update step. Using a small momentum parameter on a relatively broad distribution is likely to result in not exploring the full experimental distribution, whereas a momentum parameter that is sufficient will explore the entirety of the distribution in a manner proportional to the experimental probability distribution, visualized in Figure S9B. The effect of changing this parameter is visualized in Figure S9B.

The toy model simulations used here were performed for 50 iterations and the model distribution widths were fixed with $\sigma$ = 2.5Å. A bimodal pseudo experimental distribution was used to demonstrate the optimization of parameters increasing conformational exploration performance.

| | *A* | τ (ps) | *Tolerance* (nm) | Production time (ns) |
|---|---|---|---|---|
| β2AR modelling | 100 | 1000 | 0.1 | 5000 |
| AT1R modelling | 200 | 1000 | 0.1 | 10000 |

**Table S1**: Parameters used in the modelling pipeline here

| Comparison | Statistic | p-value | Lower CI | Upper CI |
|---|---|---|---|---|
| 1-2 | -3.536 | <1E-10 | -4.317 | -2.755 |
| 1-3 | 3.603 | <1E-10 | 2.822 | 4.384 |
| 1-4 | 1.778 | 9.48E-08 | 0.997 | 2.559 |
| 2-1 | 3.536 | <1E-10 | 2.755 | 4.317 |
| 2-3 | 7.139 | <1E-10 | 6.358 | 7.92 |
| 2-4 | 5.314 | <1E-10 | 4.533 | 6.095 |
| 3-1 | -3.603 | <1E-10 | -4.384 | -2.822 |
| 3-2 | -7.139 | <1E-10 | -7.92 | -6.358 |
| 3-4 | -1.825 | 4.23E-08 | -2.606 | -1.044 |
| 4-1 | -1.778 | 9.48E-08 | -2.559 | -0.997 |
| 4-2 | -5.314 | <1E-10 | -6.095 | -4.533 |
| 4-3 | 1.825 | 4.23E-08 | 1.044 | 2.606 |

Table S2: Tukey's HSD test of L70-P299 Cα distances between each generated conformation ensemble. The top 50 models in each conformation were used (displayed in Figure 5B). The test statistic, computed for each pairing, is the difference between the sample means. 95% confidence intervals of the test statistic are also provided (Lower and Upper CI).

| Component/Site | 55 | 139 | 220 | 236 | 304 | 311 |
|---|---|---|---|---|---|---|
| 1 | 3.02 | 2.84 | 1.90 | 2.86 | 2.53 | 4.12 |
| 2 | 3.28 | 2.51 | 1.89 | 1.84 | 3.64 | 5.63 |
| 3 | 1.74 | 2.39 | 1.54 | 2.24 | 2.59 | 3.56 |
| 4 | 2.15 | 3.19 | 2.38 | 1.71 | 2.44 | 4.68 |

Table S3: RMSF values of spin site Cα atoms for the top 50 models in each conformation. Components are listed as row indices and spin sites are listed as column indices. Values in the table are in angstroms (Å).

| Comparison | Statistic | p-value | Lower CI | Upper CI |
|---|---|---|---|---|
| 1-2 | -2.404 | <1E-10 | -3.265 | -1.543 |
| 1-3 | -0.008 | 1.000 | -0.869 | 0.853 |
| 1-4 | -1.445 | 1.29E-04 | -2.306 | -0.584 |
| 2-1 | 2.404 | <1E-10 | 1.543 | 3.265 |
| 2-3 | 2.396 | <1E-10 | 1.535 | 3.257 |
| 2-4 | 0.959 | 0.022 | 0.099 | 1.82 |
| 3-1 | 0.008 | 1.000 | -0.853 | 0.869 |
| 3-2 | -2.396 | <1E-10 | -3.257 | -1.535 |
| 3-4 | -1.437 | 1.42E-04 | -2.298 | -0.576 |
| 4-1 | 1.445 | 1.29E-04 | 0.584 | 2.306 |
| 4-2 | -0.959 | 0.022 | -1.82 | -0.099 |
| 4-3 | 1.437 | 1.42E-04 | 0.576 | 2.298 |

Table S4: Tukey's HSD test of D125 Cγ-R126 Cζ distances between each generated conformation ensemble. The top 50 models in each conformation were used (displayed in Figure 5B). The test statistic, computed for each pairing, is the difference between the sample means. 95% confidence intervals of the test statistic are also provided (Lower and Upper CI).

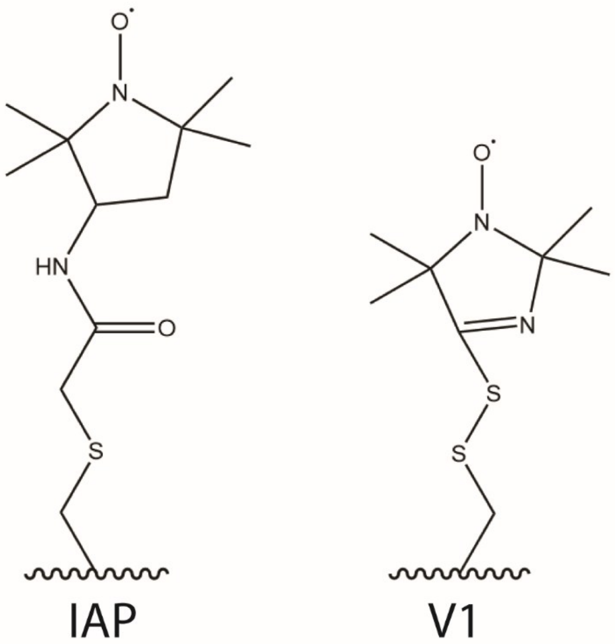


Figure S1: IAP (left) and V1 (right) spin labels. Curved lines represent the backbone of the protein.

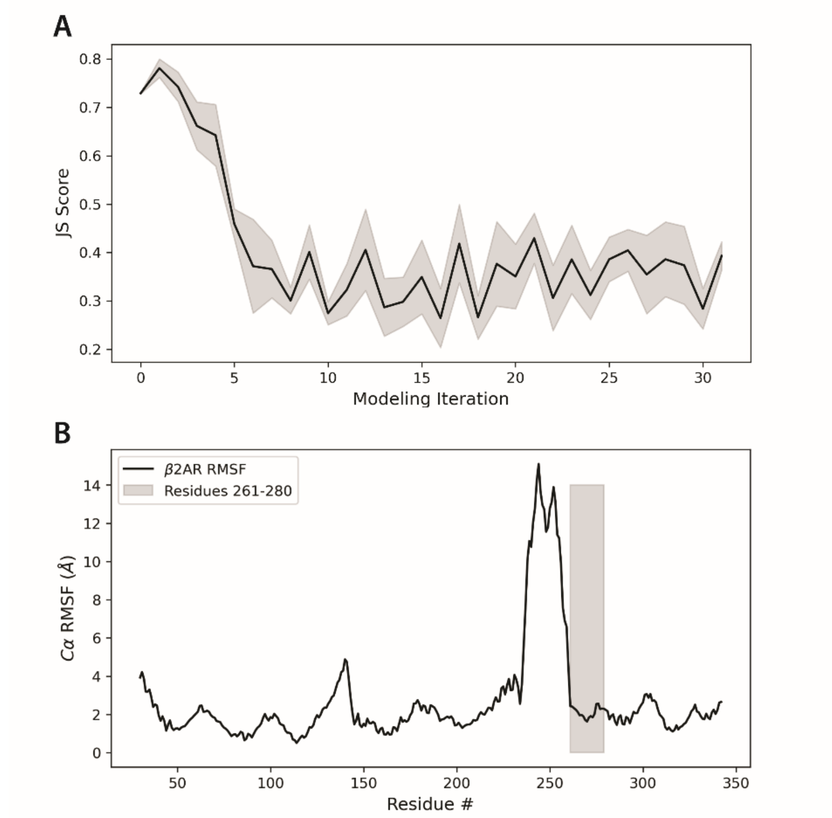


Figure S2: A) JS divergence improvement across iterations. The average JS divergence across all replicate modelling iterations was plotted with the SEM shown (grey). B) Cα RMSF of the top 10 β2AR models. The grey-shaded region indicates residues 261-280 of TM6.

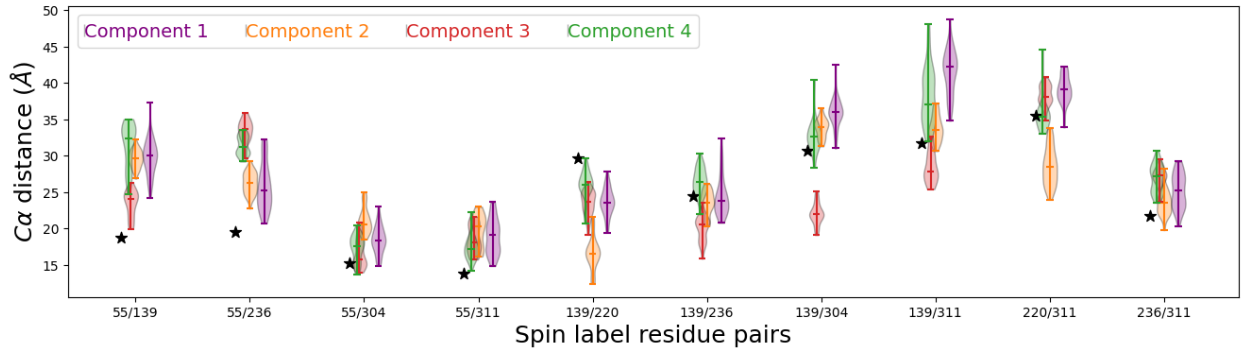


Figure S3: Violin plot of Cα distance between spin label residue pairs for the top 50 models generated from each NMF component. The body width of each violin represents the density of points, the middle bar represents the median value, and the max and minimum bars represent the extrema of the Cα distance values. Black stars represent the Cα distance between spin label residue pairs of the starting model.

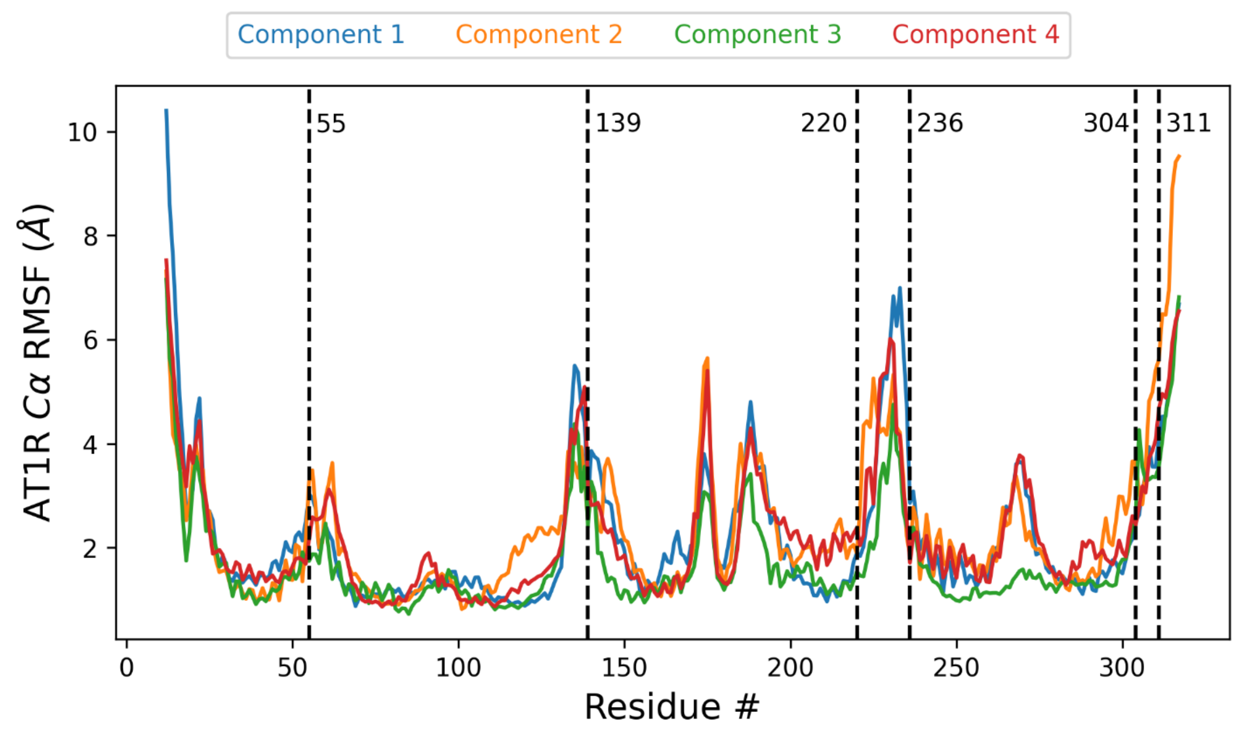


Figure S4: RMSF plot of top 50 models for each AT1R conformation. Dashed black lines represent Cα atoms of spin label sites with the specific site labeled next to each line.

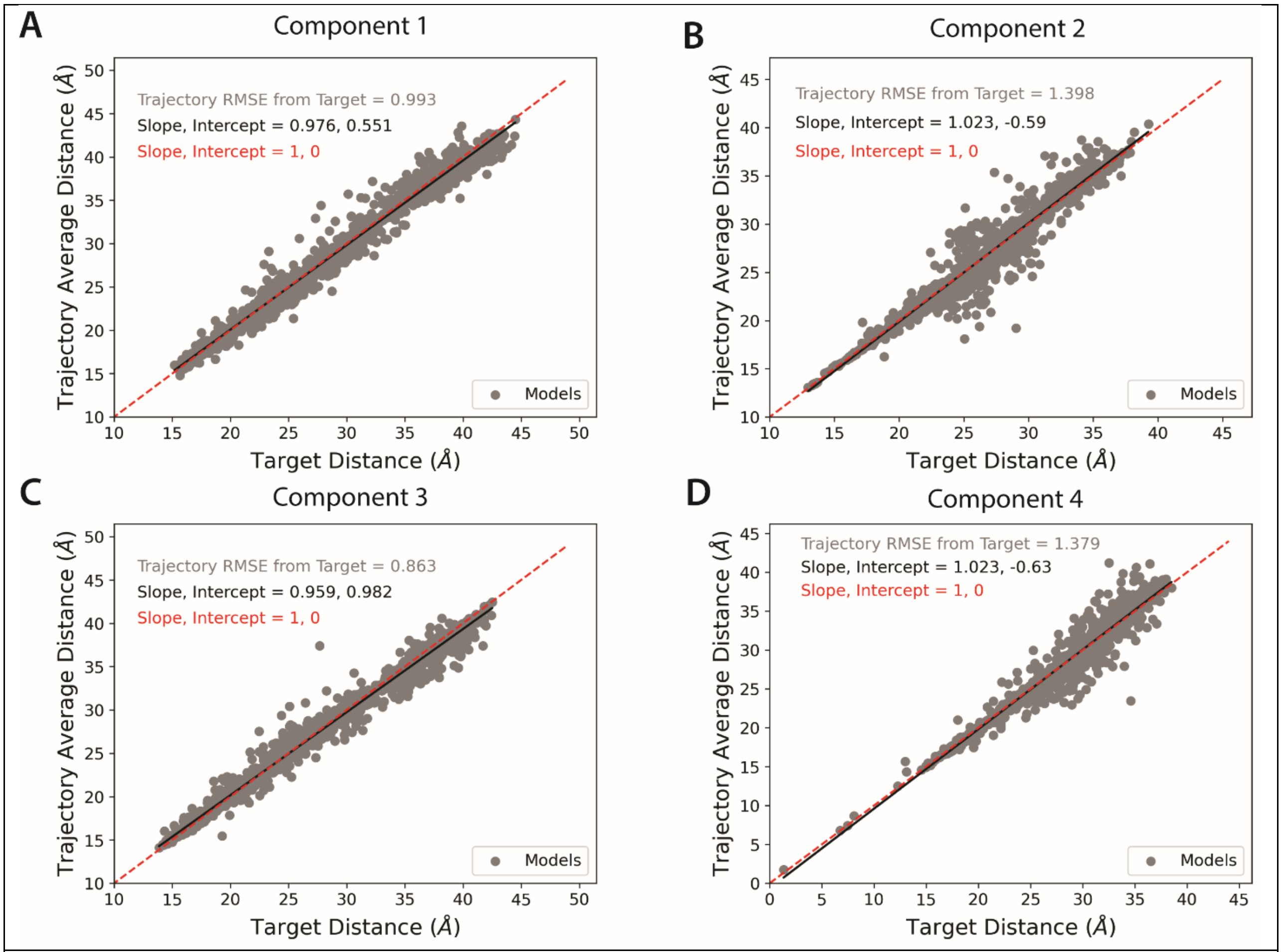


Figure S5: Average trajectory Cα spin pair distance vs. target distance for components 1-4 (A-D, respectively). Models and their Cα distances for each site are represented by grey dots. A line of fit to the data is shown as a black line with the corresponding slope and intercept. The diagonal for the plot is shown as a red dashed line for comparison with the line of fit.

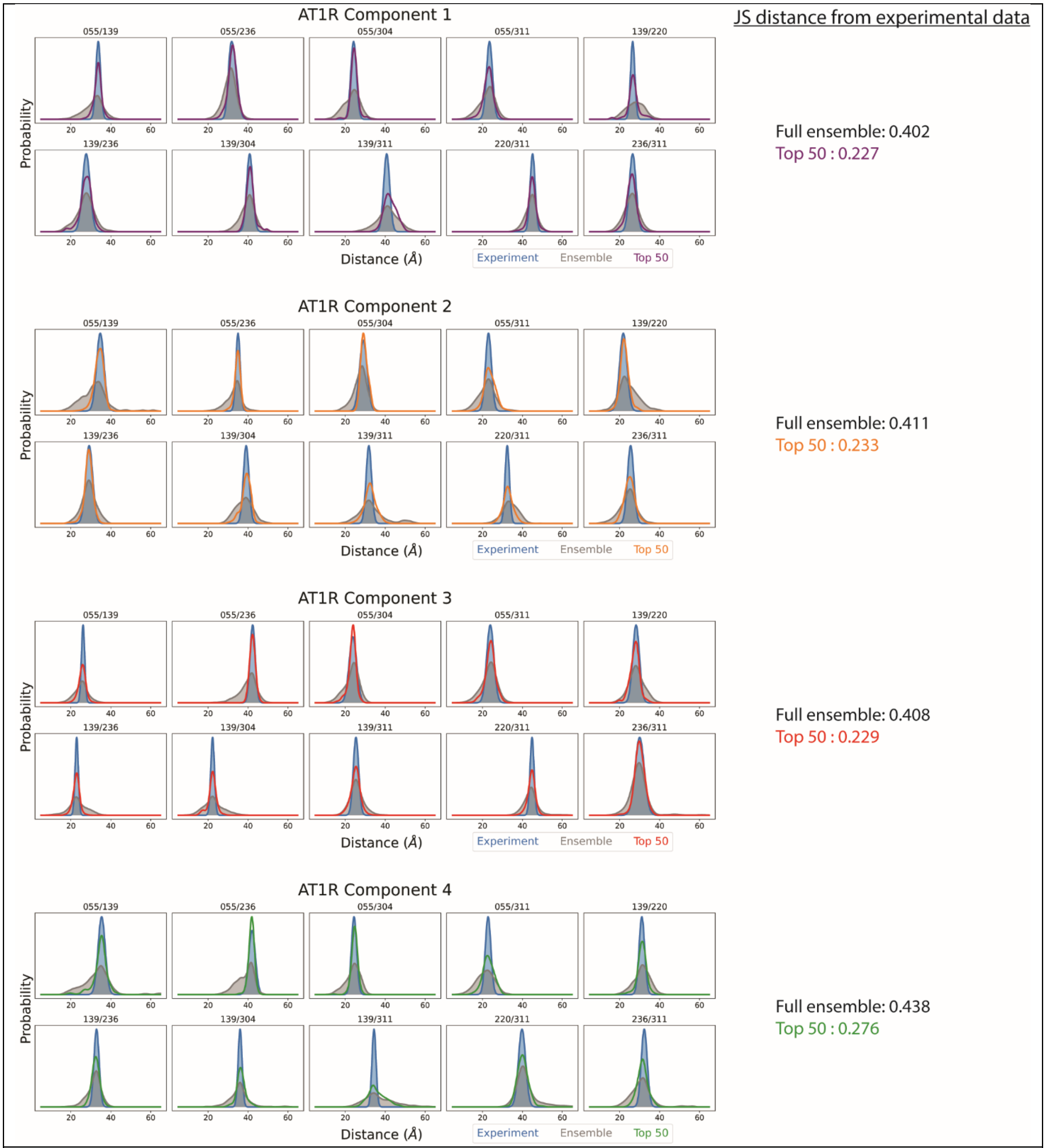


Figure S6: JS divergence from experimental data of the full model ensemble and top 50 models of each conformation.

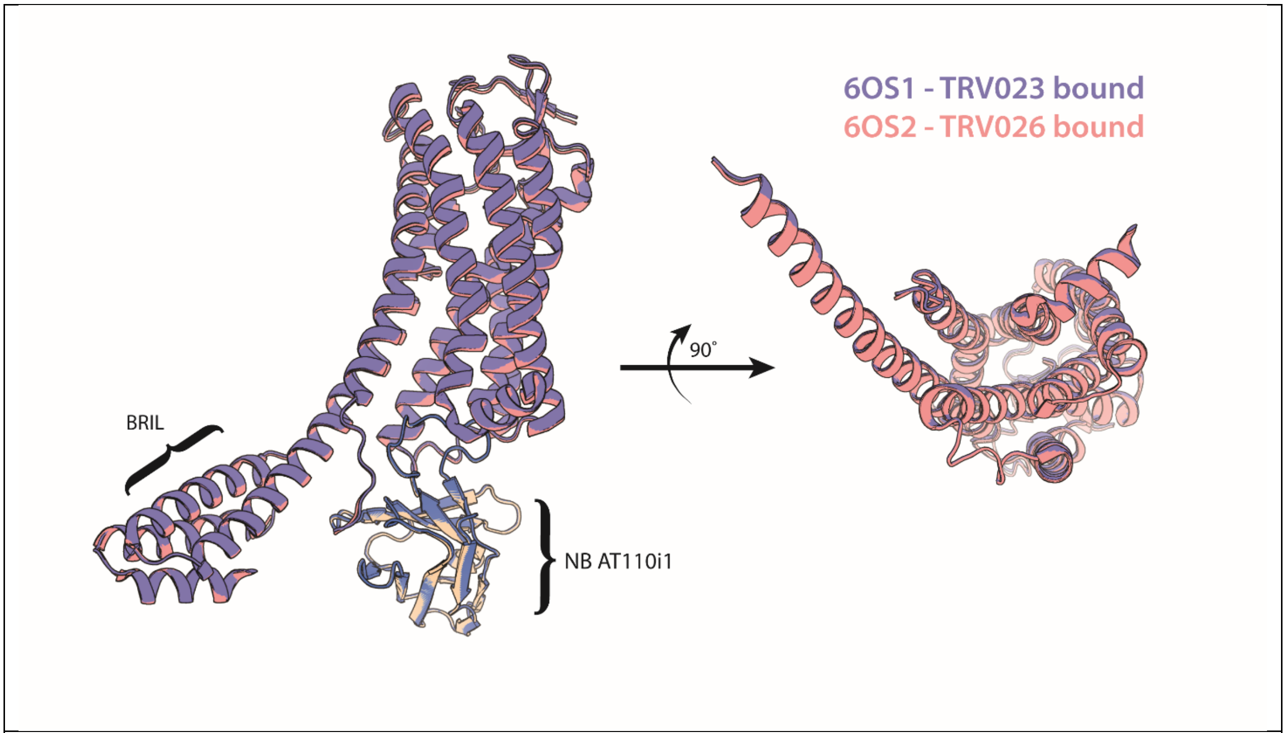


Figure S7: Overlay of X-ray crystal structures 6OS1 and 6OS2. Added BRIL and bound NB AT110i1.

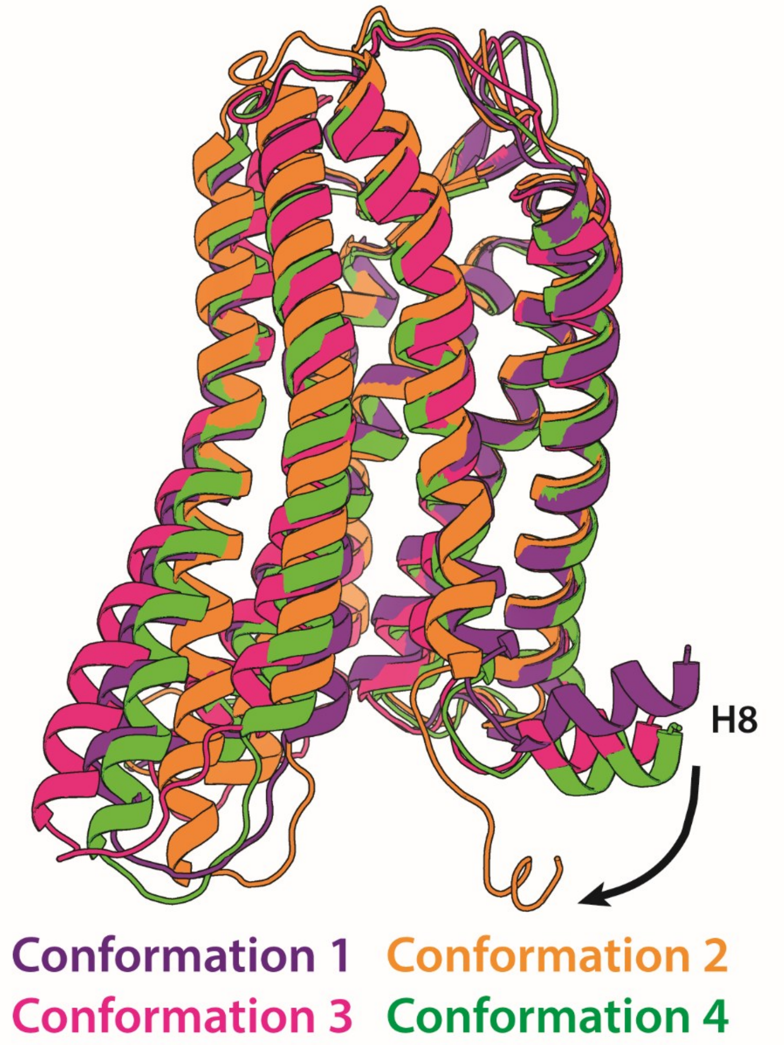


Figure S8: Average models from all conformations showing the significant rearrangement of H8 in conformation 2.

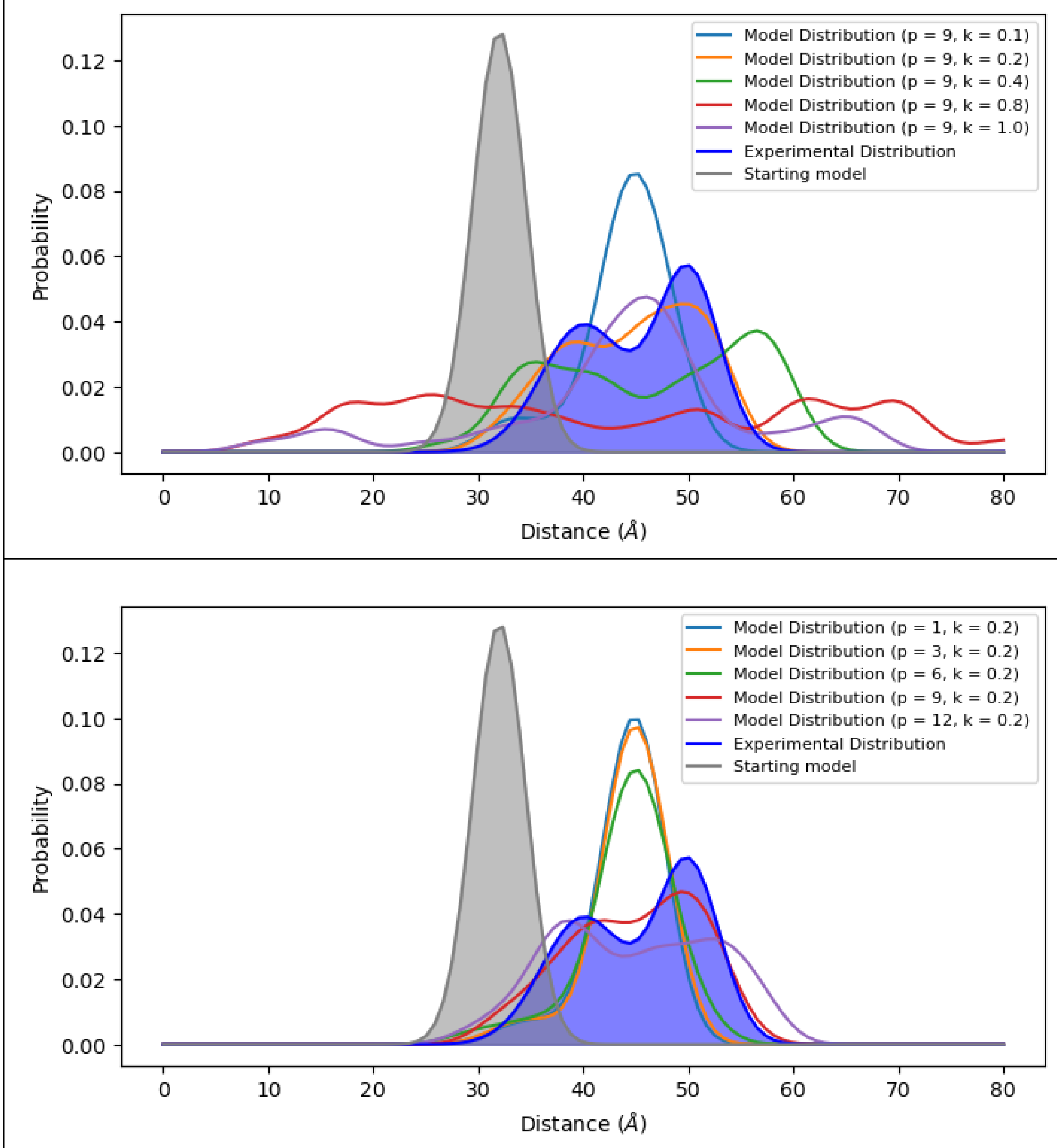


Figure S9: Effect of model parameters on modeling algorithm performance using a toy-model system of two points on a line. Starting model distributions are shown in grey and experimental distributions are shown in blue. 50 iterations were carried out and cumulatively summed to show the overall Model Distribution. Varying learning rate (k) values (top) and momentum (ρ) values (bottom) and their effects on the toy-model system exploration are graphed here. Small values of ρ may not sample the entire experimental distribution and large values of ρ may oversample the experimental distribution. Similarly, small values of k may not sample the entire experimental distribution and large values of k lead to system instability. These parameters can be tuned for each molecular system and experimental data set to produce modeled distributions (without refinement) similar to those represented by the ρ=9/k=0.2 parameter distributions here, even in bimodal distance distributions.
